# Reducing RSV hospitalisation in a lower-income country by vaccinating mothers-to-be and their households

**DOI:** 10.1101/569335

**Authors:** Samuel P. C. Brand, Patrick Munywoki, David Walumbe, Matt J. Keeling, D. James Nokes

## Abstract

Respiratory syncytial virus is the leading cause of lower respiratory tract infection among infants. RSV is a priority for vaccine development. In this study, we investigate the potential effectiveness of a two-vaccine strategy aimed at mothers-to-be, thereby boosting maternally acquired antibodies of infants, and their household cohabitants, further cocooning infants against infection. We use a dynamic RSV transmission model which captures transmission both within households and communities, adapted to the changing demographics and RSV seasonality of a low-income country. Model parameters were inferred from past RSV hospitalisations, and forecasts made over a 10-year horizon. We find that a 50% reduction in RSV hospitalisations is possible if the maternal vaccine effectiveness can achieve 75 days of additional protection for newborns combined with a 75% coverage of their birth household co-inhabitants (∼7.5% population coverage).

## Introduction

Respiratory syncytial virus (RSV) is the most common viral cause of acute lower respiratory infection ***Nair et al.*** (***2010***). A large majority of children contract RSV by the age of two ***Glezen et al.*** (***1986***); ***Ohuma et al.*** (***2012***) but the chance of developing severe disease from a RSV infection is much greater amongst young infants (<6 months) ***Hall et al.*** (***2009***) and decreases rapidly with the age of the infected. Vaccine development aimed at protecting young children against RSV disease has become a global health priority ***World Health Organization*** (***2017***). As of December 2018 there are over 40 RSV vaccines in development ***PATH*** (***2018***). In particular, two vaccination approaches have been identified as potentially effective: a single dose vaccine aimed at mothers-to-be leading to antibody transfer across the placenta thereby boosting maternally acquired immunity among newborns, and paediatric vaccination aimed directly at infants ***Modjarrad et al.*** (***2016***); ***World Health Organization*** (***2017***). Moreover, it is possible that a prophylactic extended half-life monoclonal antibody could act as a vaccine surrogate whilst replicating the desired effect of a maternal vaccine ***Zhu et al.*** (***2017***); ***Domachowske et al.*** (***2018***). A serious complication in RSV vaccine development has historically been the risk of causing enhanced disease amongst the immunologically naive ***Chin et al.***(***1969***), therefore it might be more prudent to target a paediatric vaccine at older children with better developed immune systems rather than young infants most at risk of RSV disease ***Anderson et al.*** (***2013***).

The desirous effect of vaccinating older children is two-fold: the vaccine both decreases the risk of morbidity in the vaccinated child and reduces the risk of transmission from the older child to any young infant the vaccinated child contacts ***Anderson et al.*** (***2013***). Molecular analysis of nasopharyngeal samples collected from a semi-rural community in Kenya has identified that the majority of RSV infections among young infants originated from within their household rather than the wider community, with older siblings being the usual household index case ***Munywoki et al.*** (***2014***), echoing a previous household study of RSV transmission ***Hall et al.*** (***1976***), although it should also be noted that the young infant was herself the index case on a significant number of occasions. This finding emphasises that reducing transmission to young infants within the household could be an effective way of reducing RSV disease in low- and middle-income countries (LMICs). However, the significant number of young infant index cases within households suggest that ‘cocooning’ young infants from transmission by vaccinating others in their household may not be sufficient by itself. Ideally, cocoon protection should be achieved in conjunction with directly protecting the young infants using a maternal vaccine.

In this paper we assess the efficacy of a mixed vaccination strategy in a LMIC setting, Kilifi county Kenya. In our scenarios there were at least one maternal vaccine and one paediatric vaccine available as per WHO priority ***World Health Organization*** (***2017***). In Kenya there are very high rates of antenatal contact between pregnant women and health professionals (97.5% in Kilifi county; ***KNBS*** (***2015***)). This suggested targeting pregnant women as part of their antenatal contact, and then offering the paediatric vaccine to all over one year olds, including adults, cohabiting with the pregnant mother. The essential idea was to leverage antenatal contact to achieve a very high coverage of a maternal antibody boosting (MAB) vaccine, and also to target her household cohabitants with an immune response provoking (IRP) vaccine that provoked a period of immunity to RSV similar to that of a natural infection. There is evidence that a reduction in susceptibility in a vaccinated person as if a natural infection had occurred could be generic to multiple vaccination types ***Yamin et al.*** (***2016***).

Predictions of vaccine effect are derived from a dynamic transmission model designed to capture the demographic structure of the population, the seasonality of RSV transmission and how rapidly, and to whom, RSV is transmitted in both households and the wider community. Unknown model parameters were inferred using data from the large-scale long-running Kilifi Health and Demographic Surveillance System (KDHSS; ***Scott et al.*** (***2012***)), and hospitalisation admissions at Kilifi county hospital (KCH) confirmed as due to RSV since 2002. It should be noted that targeting vaccination in this way is not an approach that one would expect to greatly reduce RSV infections under the assumptions of simple compartmental models of RSV transmission because the rate of vaccination deployment would be too low (see Box 1). However, we shall see that these vaccines are efficiently targeted at creating protection for the young infants most at risk of hospitalisation if they caught RSV.

The modelling approach used in this paper differs from the majority of RSV modelling approaches extant in the literature, which largely focus on deterministic age structured transmission models ***Pitzer et al.*** (***2015***); ***Kinyanjui et al.*** (***2015***); ***Yamin et al.*** (***2016***); ***Hogan et al.*** (***2016***), although one modelling study does consider the effect of social structure on transmission using a non-seasonal approximation within a stochastic individual-based model (IBM) ***Poletti et al.*** (***2015***). Two possible explanations for the lack of household structure in RSV modelling are: first, accounting for the interplay of demography and household structure remains a significant modelling challenge ***Glass et al.*** (***2011***); ***Geard et al.*** (***2015***), and second, the dynamics of age structured transmission models can be predicted using a comparatively small set of deterministic rate equations ***Keeling and Rohani*** (***2008***). Stochastic IBMs for epidemics benefit from additional realism and flexibility compared to deterministic models. However, rigorous inference of model parameters for stochastic IBMs of epidemics is highly challenging because, along with other difficulties, the random infection times of each case will not typically be known ***O’Neill and Roberts*** (***1999***). The model used in this paper required a rate equation for each possible household configuration ***House and Keeling*** (***2008b***). Specifically for RSV modelling it has been noted that this could lead to thousands of rate equations that must be simulated simultaneously ***Kinyanjui*** (***2014***), effectively rendering the model impractical for regression against data due to slow integration time. Nonetheless, this work demonstrates that by making appropriate simplifications, and using numerical solvers adapted to large systems (in this case ∼2000 variables), it was possible to both include realistic household structure and rigorously infer model parameters for a model of RSV transmission in a LMIC setting.

## Results

The RSV transmission model parameters were either drawn from the RSV literature or inferred from age-stratified weekly hospitalisations at Kilifi county hospital (KCH) between 2002-2016. The underlying biology of the transmission model was similar to a simple compartmental model of RSV infection and waning immunity (see Box 1) with two main differences: (i) the age of the individuals affected their susceptibility to RSV, infectiousness after contracting RSV, duration of RSV infectiousness, and likelihood of developing severe disease and being hospitalised after contracting RSV, and (ii) infectious contacts were distributed at two-levels of social mixing differentiating between persistent contacts between household co-occupants and randomly assigned contacts within the community of Kilifi county based on the ages of the infected and infectee (Fig. 1 and *Methods*). The joint age and household distribution of the population accessing KCH was chosen to match the ongoing findings of the Kilifi Health and Demographic surveillance system (KDHSS; ***Scott et al.*** (***2012***)). The seasonality of RSV hospitalisations at KCH has historically been erratic with peak months for RSV hospitalisation varying as widely as November to April (appendix 1). Moreover, over the 15 year period we are studying in this paper, there was demographic change in the underlying population both in age profile and household size distribution. We addressed these modelling challenges: first, by rejecting the typical epidemiological modelling assumption that population demographic structure is at equilibrium in favour of directly modelling demographic change, and second, by treating the shifting seasonality of RSV transmission in Kilifi as being driven by an underlying latent random process to be jointly inferred with model parameters. The goal was to account for factors influencing the rate of hospitalisations that changed over the 15 years of study so as to get an unbiased estimate of parameters we assumed were static over the period, such as the person-to-person rate of transmission within a household. We were able to capture the year-to-year variation in hospitalisation, and age profile of the hospitalised, with only six free parameters (Fig. 2, *Methods*, and appendix 1). We were unable to jointly identify the rate of school children contacting other school children with the rate of homogeneous contact among all over one year olds, therefore we considered a range of within school contact rates, and for each value inferred the other six free model parameters and assessed the efficacy of vaccination for a range of MAB vaccine effectiveness values and IRP vaccine coverage values. Each scenario gave similar results for the efficacy of household targeted vaccination (see *Supporting Information*), therefore we have only presented results in the main *Results* section for the scenario with the highest rate of within school mixing. At KCH all RSV hospitalisations occurred in the under five year olds with 84% of hospitalisations occurring in the under one year olds (Fig. 2 B). This finding is consistent with the much higher rates of hospitalisation per RSV infection for younger infants ***Kinyanjui et al.*** (***2015***). However, the hospitalisation time series has to also be understood in the context of dynamic RSV transmission and demographic change in the study population. A general trend of increasing hospitalisations between 2002-2009 is at least partially explained by a 16% increase in under ones in the population over that period. The rest of year-to-year variation in hospitalisation was explained by seasonal epidemic dynamics, themselves driven by shifting seasonality (Fig. 2 A; 1).

**Figure 1.**
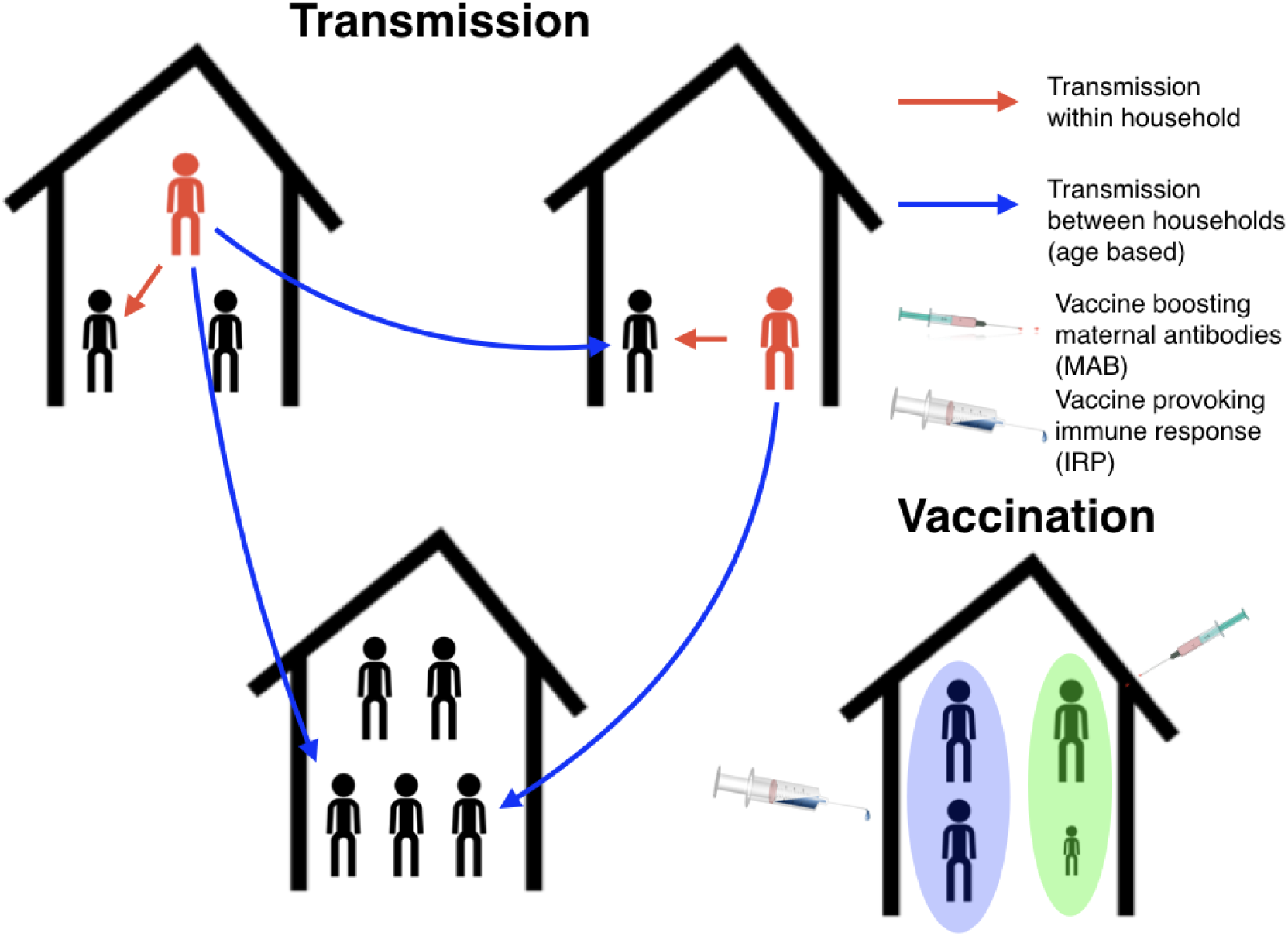
Schematic plot for the RSV transmission model and vaccination programme. Infectious individuals (red character figures) transmit to other individuals inhabiting the same house, and to other individuals in other households based on the ages of the both the infector and infectee. Red and blue arrows represent possible realised infections over a short period of time. Bottom right household demonstrates the vaccination strategy; the mother has received a maternal antibody boosting (MAB) vaccine which increased transfer of protective antibodies to newborns (green background shading), meanwhile other household members have received an immune response provoking (IRP) vaccine (blue background shading).

**Figure 2.**
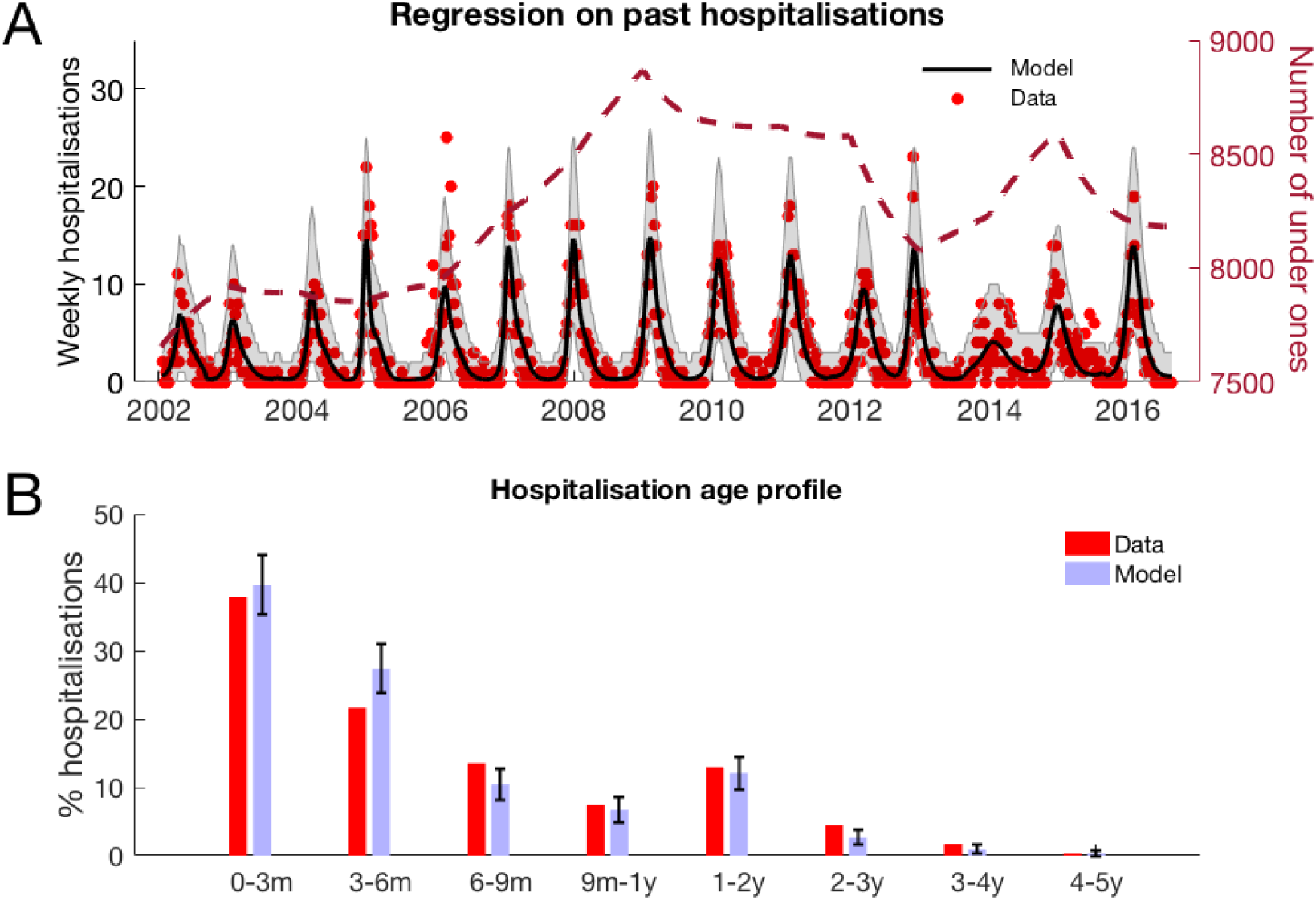
RSV hospitalisation at KCH: dynamics and age profile of hospitalised patients. **A**: Weekly RSV hospitalisations before implementation of vaccinations. Black curve gives mean prediction of RSV household transmission model after regression against weekly incidence data (red dots). Grey shaded area indicates the 99% prediction interval for the model. Also shown is the number of under ones in the population (dashed line). **B**: Age profile of hospitalisations at KCH before vaccination. Error bars give 99% prediction intervals for model.

It is not straight-forward to directly compare the reproductive ratio (*R*_0_; usually defined as the number of secondary cases created by an initial ‘typical’ case in a completely susceptible population) of transmission models with household **and** age structured contacts to those with **only** age structure ***Pellis et al.*** (***2010***). In models with only age structure *R*_0_ can be calculated from the age-based mixing rates ***Diekmann et al.*** (***1990***); the *R*_0_ for RSV in Kilifi county has been estimated as high, *R*_0_ = 7.1 or *R*_0_ = 25.6 depending on how the age-based mixing rates were estimated ***Kinyanjui et al.***(***2015***). In principle, our model did not require household transmission for RSV to persist since it would have been possible for us to infer that age-based mixing in the community dominated RSV transmission. However, if we neglect within household transmission from our model with inferred parameters we found the subcritical *R*_0_ = 0.75 for solely age-based transmission outside of the household; that is we inferred that within household transmission was necessary to sustain transmission of RSV in the population.

#### Box 1. Vaccination predictions from a simple unstructured RSV epidemic model

The essential idea in this paper is to use antenatal contact between mothers-to-be and health professionals to deploy two separate vaccines: first, a vaccine targeting the mothers-to-be which boosts the duration of protection her newborn will have against RSV (MAB vaccine), and second, a vaccine aimed at the mothers-to-be’s household cohabitants giving each a period of RSV immunity, equivalent to that of a natural infection (IRP vaccine). As a baseline for understanding RSV transmission we can use a simple mechanistic model which captures the essential biology of RSV infection; newborns are born with a period of immunity to RSV infection which is lost during their first year of life, after contracting RSV the individual is infectious for a period before gaining temporary waning immunity to RSV re-infection. Assuming homogeneous transmission the dynamics of the simple RSV transmission model can be described using four dynamic variables describing the numbers of currently maternally protected individuals (**M**), susceptibles (**S**), infecteds (**I**) and immune/recovereds (**R**). The evolution of the epidemic, after vaccination, can be given as a standard ODE:

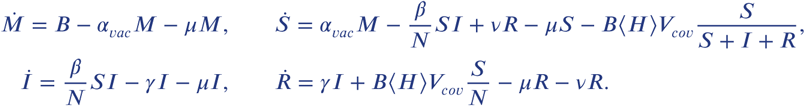

Where each term above describes the rate of events that change the epidemic state: Births (*B*), loss of maternally derived protection, after MAB vaccination, (*α*_*vac*_), mortality (*μ*), RSV force of infection (*βSI*/*N*), recovery (*γ*), reversion to susceptibility (*v*), as standard in the literature ***Anderson and May*** (***1992***); ***Keeling and Rohani*** (***2008***). The rate at which IRP vaccines successfully vaccinate susceptibles is *B ⟨ H ⟩ V*_*cov*_*S*/(*S* + *I* + *R*); that is the mean size of a pregnant woman’s household (⟨ *H* ⟩) times the effective coverage of the vaccine (0 ≤ *V*_*cov*_ *≤* 1) time the likelihood of selecting a susceptible and not wasting the vaccine assuming that we are only targeting those who have definitely lost their maternal protection to RSV (*S*/(*S* + *I* + *R*)). For simplicity, we can treat the duration of maternal protection as very short compared to the typical person’s lifetime (i.e. *α*_*vac*_ *≫ μ*). The simple RSV model is analytically tractable:

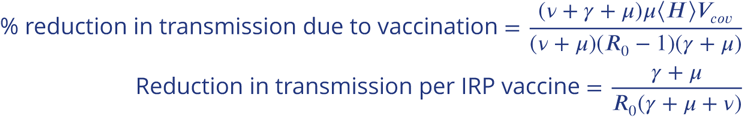

Where *R*_0_ = *β*/(*γ* + *μ*) is the reproductive ratio of RSV, and we are assuming that the birth rate is at replacement *B* = *μN*. The simple RSV model makes some general predictions about the efficacy of IRP vaccination:

- The MAB vaccine does not significantly effect transmission in the general population.
- The efficiency of the IRP vaccine (avoided infections per effective dose) should not change with coverage.
- Using parameters typical of the study population at Kilifi (see *Supporting Information*), the reduction in RSV transmission due to IRP vaccination can be modest because the deployment rate is too low; for *R*_0_ = 2 the maximum achievable reduction in transmission is *<* 4%.

Therefore, a naive simple model of RSV transmission is pessimistic about the joint vaccination strategy. However, in this study we also account for more detailed social structure, differential susceptibility, infectiousness, and risk of disease dependent on the age of the individual and seasonality in transmission. We will see that targeting vaccines socially close to young infants is much more effective than the simple model predicts.

We found that, pre-vaccination, school age children suffered on average the highest force of infection, that is the per-capita rate of infectious contacts, from outside of the household followed by under one year olds (Fig. 3 A). This finding was dependent on assuming that we had a high degree of homophily in the social contacts of school-age children (the high within school transmission scenario mentioned above). Other scenarios were considered with lower levels of in-group preference for school-age children to contact other school-age children; in the alternate scenarios the parameter imputation process found slightly higher rates of contacts within the household and homogeneously outside of the household but lead to very similar results (*Supporting Information*). The infectious contacts outside the household were distributed predominantly to individuals within households of size 2-5 (Fig. 3). This reflected the household distribution of the population; school children and under ones who were most at risk of making social contact with those infected with RSV outside the household tended to live in households of this size (Fig. 3 B).

**Figure 3.**
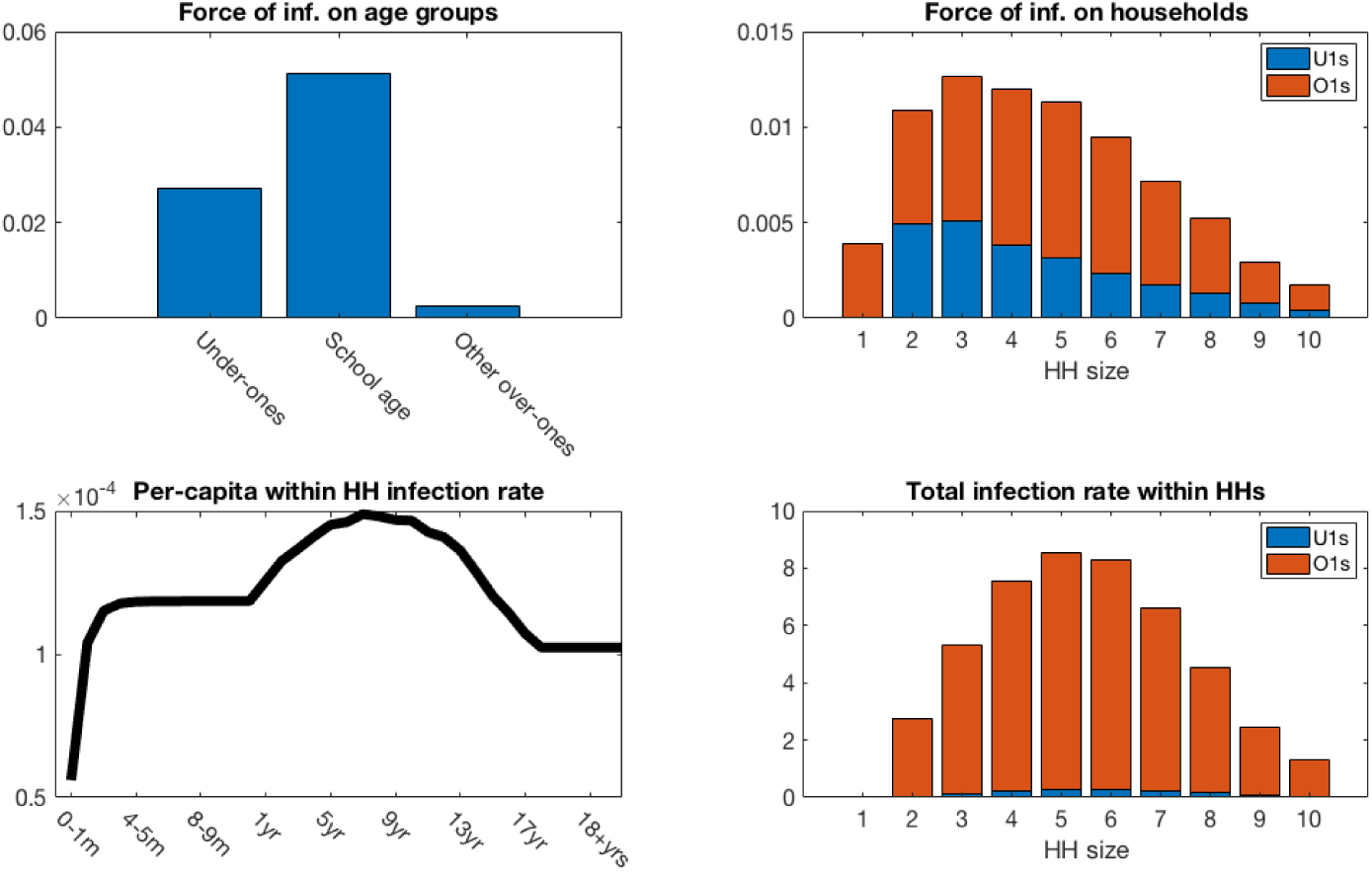
Mean force of infection (2002-2016) between households and mean infection rates within households. **A:** The mean force of infection (infectious contacts received per person per day) of RSV due to transmission from without the household on three age groups: under-ones, school age children and everyone else, including adults. **B:** Mean force of infection due to transmission without the household on individuals inhabiting each household size. **C:** The mean per-capita daily rate at which different age groups become infected with RSV from within their household. **D:** The mean total daily rate of RSV infection within households of different sizes.

Force of infection is a less natural concept for measuring within household infection due to small numbers of individuals per household, and intense frequent contacts. Instead, we measured the true rate of RSV transmission between individuals cohabiting a household. The highest per-capita rates of infection within households were for 7 year olds (Fig. 3 C); this reflected the typical age of individuals within the households most at risk of RSV introduction and with severest transmission rates after introduction. The infection rate among under ones increased rapidly until it plateaued at ∼6 months old. The rapid increase in per-capita infection rate was due to waning of maternally acquired immunity to RSV, which we inferred as lasting on average 21.6 days ([17.2, 26.1] 95% CI). The total infection rate within households was greatest in size 5 and 6 households (Fig. 3 D). This differed from the household size where each person was at most risk of contracting RSV outside the household. Two factors shifted the burden of RSV infection to larger households: first, there are more people in larger households therefore risk of RSV introduction can be higher even if the per-person rate is lower, and second, the intensity of transmission within households is higher for larger households.

We evaluated the success of the mixed, maternal antibody boosting (MAB) plus immune reponse provoking (IRP) vaccination strategy. The high antenatal contact levels in Kilifi county suggested that vaccination coverage of mothers-to-be had the potential to be very high, especially if maternal immunisation to boost newborn immunity became an established method for a range of vaccines including influenza and Group B Streptococcus. However, an available MAB vaccine might only be effective if delivered in the third trimester of pregnancy and, whilst having at least one antenatal contact is very common for pregnant women in Kilifi county, it is not clear that antenatal contact always occurs at the relevant stage of pregnancy. Therefore, we consider both a maximalist scenario (100% MAB coverage), and a more realistic uptake (50% MAB coverage). The number of days of additional maternally derived protection donated to the newborns by MAB vaccinated mothers was uncertain, we considered a range of MAB protection 0 - 90 days. We assumed that if the pregnant mother’s household cohabitants agreed to receive an immune response provoking vaccine then all were vaccinated. As is common in vaccine strategy analysis we combine coverage and effectiveness into one effective coverage (coverage times effectiveness c.f. ***Keeling and Rohani*** (***2008***)), although in this case effective coverage could be considered both within and between households. This allowed us to present our results in two main axes of vaccination unknowns: the effectiveness of the MAB vaccine (days of protection to RSV infection from birth additional to natural protection), and the effective coverage of the IRP vaccine (0-100%). We assumed that the maximum coverage of the vaccine would be reached within a year, and considered ten years of transmission after this implementation and treated the demography of the KDHSS as being unchanged from 2016 over this period. The model inference stage included inferring the statistics of yearly variation in RSV seasonality. The decrease in rates of RSV hospitalisation and infection due to vaccination over ten years presented are median improvements over 500 independent realisations of random future seasonal patterns compared to a baseline of no intervention. If the MAB vaccine was unavailable or ineffective (0 days MAB protection), we found that it was still possible to reduce RSV hospitalisations by up to 25% using only the IRP vaccine on the household members of young infants at time of birth (Fig. 4 A and B). If 100% maternal vaccination could be achieved then the MAB vaccine was more successful as a sole vaccine option compared to IRP vaccination; in the sense that 90 days of additional protection from RSV delivered a 45% reduction in hospitalisation even with no IRP vaccine coverage. Nonetheless, even with an effective MAB vaccine there was added benefit to also using a IRP vaccine; a greater than 50% reduction in hospitalisations was achieved with a MAB vaccine that gave 75 additional days of RSV protection and a 75% coverage of the pregnant womens’ households (Fig. 4 A). If only 50% maternal vaccination coverage could be achieved then unsurprisingly also using the IRP vaccine became relatively more important. The mixed vaccination strategy that achieved better than 50% hospitalisation reduction with 100% maternal coverage achieved 38% reduction in hospitalisations with 50% maternal coverage (Fig. 4 B); reducing the maternal coverage by half didn’t necessarily half the success of the vaccination programme so long as IRP vaccine was also available. Improving the effectiveness of the MAB vaccine caused a significant improvement in hospitalisations, but had an almost negligible effect on the total infections in the population (Fig. 4 C and D). IRP vaccination was more effective at reducing total RSV infections, but even at 75% coverage of the households of women giving birth the reduction in infections was *<* 4% (Fig. 4 C and D). That IRP vaccination had a modest effect on the true infection rate, and that MAB vaccination has a negligible effect on the true infection rate, was in line with the prediction of the simple non-seasonal RSV model (Box 1). However, the simple model could not predict that the percentage reduction in hospitalisations would be significantly greater than for total infections because of the direct and indirect protection of those most at risk of disease. For the mixed strategy achieving a 50% reduction in RSV hospitalisations at 100% MAB coverage described above the seasonal dynamics of hospitalisations post-vaccination were similar to “no intervention” but at a lower rate (Fig. 5 A). There was a reduction in median hospitalisations in every age group, but predominantly in 0-3 month years old (who are nearly all protected by the MAB vaccine) and 3-6 month year olds (Fig. 5 B). However, targeting pregnant women and their cohabitants did not prevent sufficient RSV infections as to significantly disrupt RSV transmission within the population at large. Nonetheless, those who were protected were overwhelmingly among those at most risk of disease if they had caught RSV.

**Figure 4.**
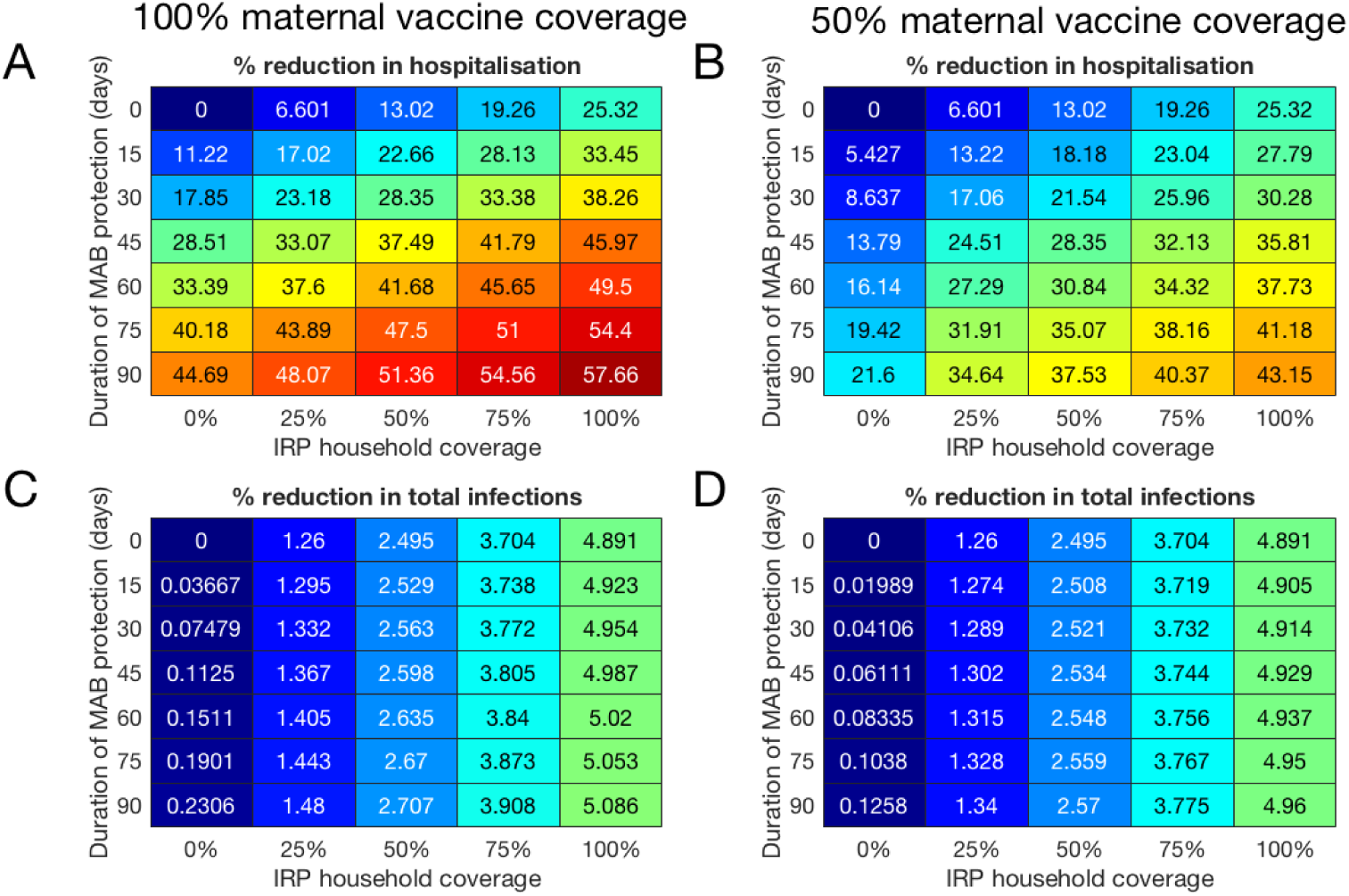
Forecast effectiveness of RSV vaccination for different mixed strategies over a 10 year period for 100% maternal vaccine effective coverage (**A** and **C**) and 50% maternal vaccine effective coverage (**B** and **D**). **A** and **B:** Percentage reduction in hospitalisations at KCH. **C** and **D:** Percentage reduction in total RSV infections in the population.

**Figure 5.**
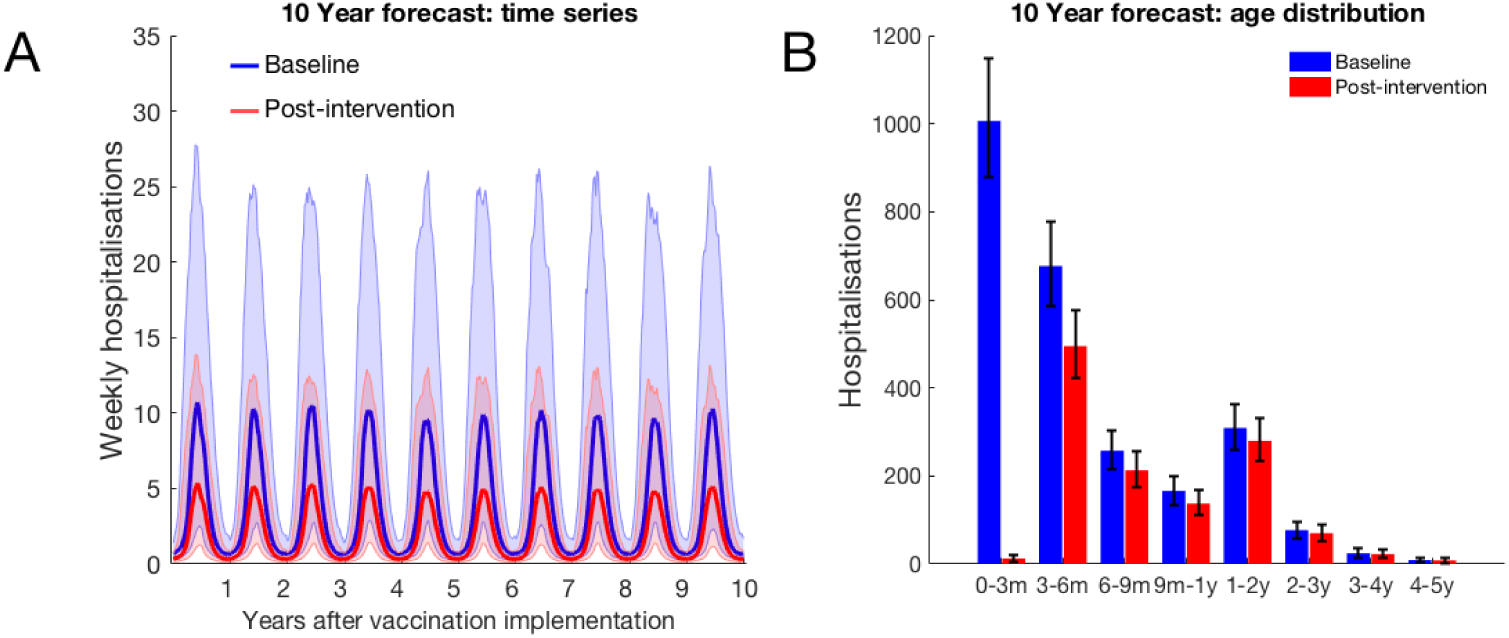
10 year forecast of RSV vaccination effectiveness for a mixed strategy of an MAB vaccine provided 75 days of additional RSV protection for newborns and a 75% IRP vaccine household coverage. **A:** Forecast weekly hospitalisations for a baseline of no vaccination (*blue*) and the mixed vaccination strategy (*red*). Shown are median forecast (*curves*) and 95% prediction intervals (*background shading*). **B:** Forecast age distribution of total RSV hospitalisations at KCH. Median forecast (*bars*) and 95% prediction intervals (*error bars*).

Over 10 years of forecasted RSV epidemics if a MAB vaccine was available, and given to every pregnant mother, 8,601 MAB vaccines were deployed each year, 0 otherwise. 0 - 24,095 *effective* IRP vaccines were deployed each year depending on effective coverage. It should be noted that by 2016 the KDHSS population was around 240,000 people, hence 100% effective coverage of the households where births occurred corresponded to ∼10% effective coverage of the total population. We measured the efficiency per vaccine with a few different metrics: both avoided hospitalisations and total infections per vaccine compared to a baseline of no-intervention as well as the extra avoided hospitalisations and total infections per effective IRP vaccine only compared to only using a MAB vaccine. Note that we are presenting results relative to *effective* coverage. In each case we considered median number of RSV hospitalisations/infections avoided. The MAB vaccine was more efficient than IRP vaccination per vaccine at avoiding hospitalisations so long as greater than 15 days of additional protection for newborns from RSV infection could be achieved. When an effective MAB vaccine was available increasing IRP vaccination decreased the efficiency per vaccine at avoiding hospitalisations from the higher MAB vaccine only efficiency towards the efficiency of IRP only vaccination (Fig. 6 A). On the other hand, IRP vaccination was more efficient at reducing infections per vaccine, conversely to the reduction in hospitalisations per vaccine the efficiency per vaccine increased towards the efficiency of IRP only vaccination (Fig. 6 B). In either case the efficiency of IRP vaccination compared to a baseline of MAB vaccination only did not change with effective coverage. IRP vaccination efficiency being independent of effective coverage is in line with what one might expect from a homogeneous mixing RSV model (see box 1).

**Figure 6.**
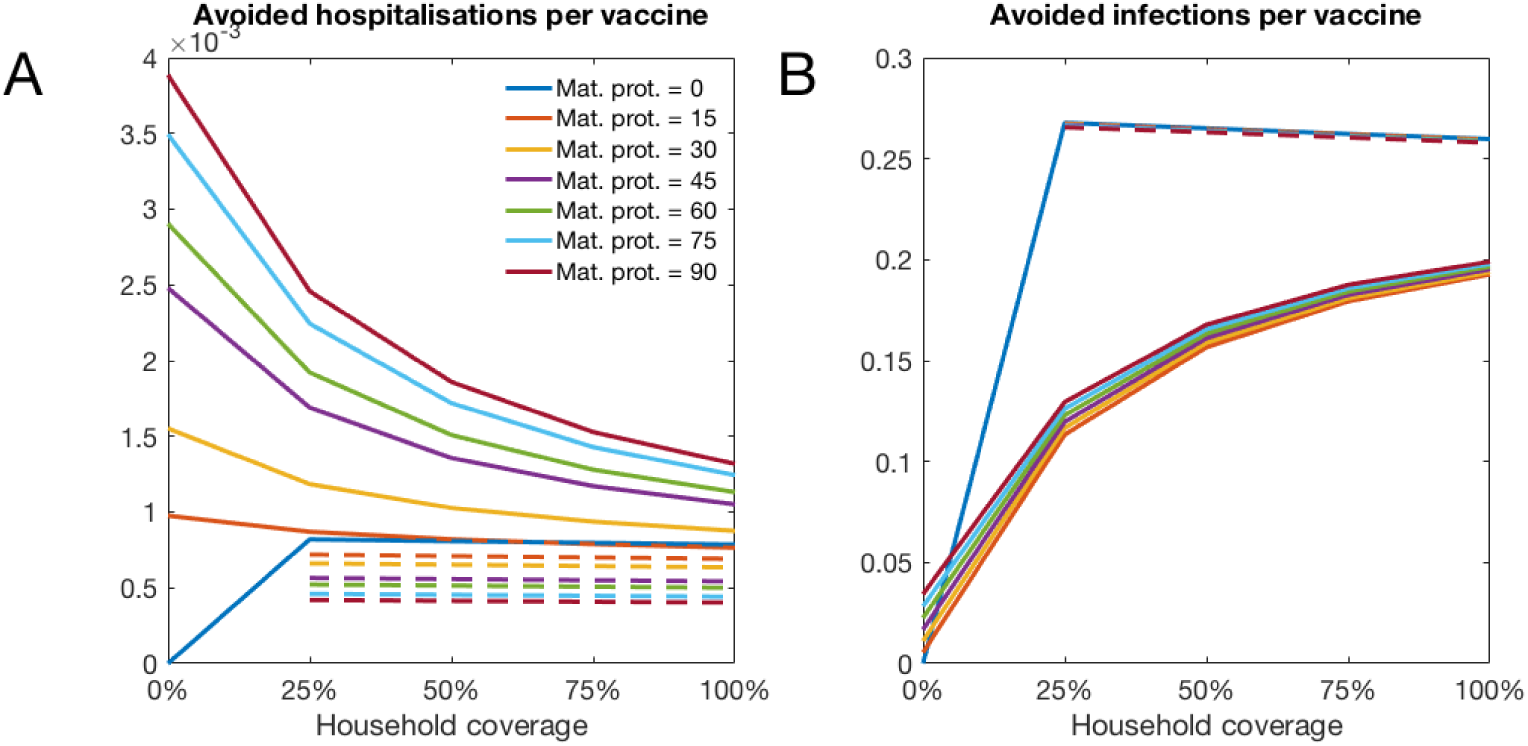
The forecast efficiency of vaccination against RSV for different mixed strategies over a 10 year period. Solid lines show the overall vaccine efficiency compared to a baseline of no intervention, dashed lines show the vaccine efficiency per IRP vaccine compared to a baseline of only MAB vaccination. **A:** Median avoided hospitalisations at KCH per vaccine over 500 simulations. **B:** Median avoided RSV infections in population per vaccine over 500 simulations.

## Discussion

Our modelling analysis suggested that a high coverage vaccination campaign of mothers-to-be with a vaccine inducing elevated levels of transplacenta RSV antibody transfer to her newborn, alongside targeting the newborn’s cohabitants with a generic vaccine that provoked a period of immunity to RSV can achieve greater than 50% reduction in hospitalisations due to RSV. This combined vaccination strategy suggested itself due to the high antenatal contact rates between mothers-to-be and health professionals in Kilifi county, Kenya (97.5% ***KNBS*** (***2015***)). We found that the combined vaccination strategy was efficient at targeting effort towards cocooning the young infants most at risk of developing RSV disease from contact with RSV infected individuals. Even at maximum effective household coverage for the IRP vaccination only ∼10% of the population were vaccinated each year with a modest reduction in the RSV infection rate of ∼5%. Nonetheless, at that coverage IRP vaccination alone achieved a 25% reduction in hospitalisations at KCH even without an effective MAB vaccine to provide direct protection to young infants. This demonstrated that although we were vaccinating at a low rate compared to population size, with only a modest reduction in infection rate, those people we did vaccinate were efficient at cocooning young infants from transmission and therefore risk of severe disease. If an effective MAB vaccine was also available the reduction in hospitalisations was greater, although the additional protection due to cocooning was relatively less since young infants were also protected from contracting RSV at the age when they were at most risk of severe disease.

We constructed the model used in this paper with the purpose of estimating the efficacy of targeting pregnant women and their households for vaccination. In order to make predictions mechanistic models of disease transmission must approximate the social structure of the population being modelled, and hence the contact rates between individuals. The focus on household transmission in this paper necessitated including households into the modelled social structure; this represented significant additional effort in model construction, computational resource and inference compared to simpler models. A more common approach in the literature is to treat the contact rates between individuals as being determined only by their respective ages. This approach has the benefit of being conceptually straight-forward and draws on a number of recent and high-quality studies which quantify social contact patterns by age stratification ***Mossong et al.*** (***2008***); ***Kiti et al.*** (***2014***); ***Prem et al.*** (***2017***). However, the fundamental theory of age-structured transmission models for endemic diseases was developed mainly with reference to diseases that induce very long term or lifelong immunity ***Anderson and May*** (***1992***). For diseases provoking long lasting immunity one would expect most older household members to be immune and therefore household structure to be a relatively less important factor in predicting risk of transmission compared to the age-structured transmission outside of the household. Indeed, simulation study of a generic strongly immunizing infection with realistic demography found limited difference in predicted incidence rate by age for people at schooling age or older between models with household structure **and** age structure compared to models with **only** age structure ***Geard et al.*** (***2015***). However, it is not clear that neglecting household structure is a good approximation for modelling seasonal RSV transmission for two reasons: first, previously infected people lose effective immunological protection to RSV rapidly enough that each season could be closer to an ’epidemic’ scenario rather than an ’endemic’ scenario. Second, every hospital admission at KCH confirmed as due to RSV was a pre-school aged child; in contrast to predicted incidence rates for school age and older individual, the simulation study cited above ***Geard et al.*** (***2015***) predicted that incidence was lower for 0-5 year olds, especially so for under one year olds, once household structure was taken into account. It would be of great interest to have a more general theoretical understanding of which epidemiological questions require household structure, or a more general meta-population structure, for epidemiological modelling, and which don’t. This remains an active area of research ***Ball et al.*** (***2015***).

A cocooning protective effect of households could explain the big discrepancy between our estimate of the mean period of protection against RSV after birth due to transplacental transfer of antibodies from mother to baby in the the womb (21.6 days of natural protection on average) compared to a RSV transmission modelling study by Kinyanjui *et al* on the same population using an age-structured model ***Kinyanjui et al.*** (***2015***) (2.3 months of natural protection if the age mixing was based on diary estimates of contacts ***Kiti et al.*** (***2014***) or 4 months of natural protection if the age mixing was based on household co-occupancy and schooling ages). The age-structured model used in the Kinyanjui *et al* study reported high or very high reproductive ratios: 7.08 for the diary based contact patterns, and 25.60 for the household co-occupancy and schooling age based contact pattern. Therefore, to fit the KCH hospitalisation data the age structured model necessarily predicted a very high level of natural protection due to maternal antibodies to compensate for the predicted high force of infection on young infants. In our model we included household structure and we fit to the same KCH data but with a much lower level of natural protection from RSV. This in turn changes the guidance modelling gives to vaccination strategy; some age structured RSV transmission models have emphasized reducing force of infection by vaccinating infants directly ***Kinyanjui et al.*** (***2015***), and 1nd that maternal vaccination is likely to be of limited impact ***Pan-Ngum et al.***(***2016***), because they have inferred that the RSV reproductive ratio is high and, therefore, natural protection to RSV is also inferred to be high. In contrast, we infer that natural protection to RSV is low and therefore 1nd that maternal vaccination in combination with elevating the cocoon protection to young infants provided by vaccinating household co-inhabitants is a highly efficient strategy. Another age-structured RSV transmission model ***Yamin et al.*** (***2016***) has found that vaccinating under-fives to RSV along with their influenza vaccination was highly efficient because of the large number of secondary cases generated per infected under-five year old. Again, it is not clear whether this result extends to a population structured into households where it is known that clustering in contacts has a complex interplay with disease dynamics, either reducing spread because infectious contacts are ‘trapped’ in the local cluster (e.g. the household) or promoting spread by enhancing persistence ***Miller*** (***2009***); ***Sun et al.*** (***2015***).

This was a modelling study and, as ever, there are factors that we have neglected in our analysis that could be addressed in future work. First, we treated coverage of the maternal vaccine and the IRP vaccine as independent. In reality, the simplest and cheapest scenario whereby the household cohabitants of pregnant mothers are recruited to the vaccination programme is if they attend antenatal contact with the mother-to-be late in the pregnancy. Therefore, if many pregnant women in their third trimester, when a MAB vaccine dose is expected to be effective, are missed from the vaccination programme then their household cohabitants would also be missed. Our results suggest that a MAB vaccine at a high coverage sharply reduces RSV hospitalisation even when the amount of additional protection is low (15 days) and if the MAB vaccination coverage is reduced to 50% IRP coverage becomes relatively more important to reducing hospitalisations. Therefore, it could be cost effective to devote extra resources towards encouraging pregnant women, and their cohabitants, to return for vaccination late in the pregnancy if their initial antenatal contact their before her third trimester. Second, the cost per vaccine remains unknown and we have not considered any measurement of the burden of disease other than hospitalisations at KCH. RSV hospitalisations have been identified as a crude proxy for the true disease burden; the passive reporting of RSV hospitalisation can vary for reasons completely independent of RSV epidemiology ***Modjarrad et al.*** (***2016***). Third, despite accounting for demographic change in our inference of model parameters we neglect demographic change in our forecasting. Our primary goal in this paper has been to establish the importance of thinking jointly about hospitalisation risk, population structure (in particular household co-occupancy) and future vaccination programmes. These issues would suggest that RSV vaccination policy would benefit from further cost-benefit analyses tailored to LMIC settings, possibly using more flexible stochastic IBMs with the model parameters inferred in this study.

In conclusion, in this paper we have analysed the performance of a joint maternal and household targeting RSV vaccination strategy measuring both reduction in hospitalisations and the true population incidence rate. We drew our conclusions based on rigorous inference of underlying transmission parameters and the inherent protection to RSV newborns received from their mothers, taking into account potential confusing factors such as variable seasonality and demography. Two central insights from our study were that the duration of natural protection to RSV that newborns inherit from their mother was likely to be much shorter than previously estimated and that RSV attack rates within the household were significant in maintaining RSV transmission. Therefore, targeting pregnant women and their households for RSV vaccination is likely to be an effective and efficient strategy under a wide range of different scenarios.

## Methods

The dynamical RSV model used in this paper simulated infection and transmission of RSV among a population described by the Kilifi Demographic and Health surveillance system (KDHSS ***Scott et al.***(***2012***)) between September 2001 to September 2016. The population was assumed to mix and transmit RSV at two social levels: within their household and outside their household among the wider community. RSV infection was modelled using a modified version of the classic susceptible, infected, recovered (SIR) compartmental framework ***Anderson and May*** (***1992***); ***Keeling and Rohani*** (***2008***). The main modifications were consistent with previous RSV transmission models; we assumed that: (i) individuals were born with a temporary immunity to RSV which faded over time, and (ii) RSV infection episodes provide individuals with only temporary protection from re-infection (mean 6 months ***Scott et al.*** (***2006***)) ***White et al.*** (***2007***); ***Moore et al.*** (***2014***); ***Pitzer et al.*** (***2015***); ***Kinyanjui et al.*** (***2015***); ***Yamin et al.*** (***2016***). The high dimensionality of the ODE model (see below) used in this paper necessitated a relatively simple compartmental structure for RSV infection progression, therefore the population is only crudely age stratified into under-one year olds (U1s) and over-one year olds (O1s). However, more detailed information about the age of the individuals in the model was available by considering their age distributions conditional on their crude age category and the type of household they inhabited (see below). After an initial RSV infection there is evidence that individuals retain reduced susceptibility to subsequent RSV infection ***Henderson et al.*** (***1979***); ***Hall et al.*** (***1991***), and will potentially have less infectious asymptomatic episodes if infected ***Hall et al.*** (***2001***); ***Yamin et al.*** (***2016***). Some RSV transmission models, using simpler social structures, therefore allow individuals to be characterised by both their age and their number of previous RSV infections ***Kinyanjui et al.*** (***2015***); ***Yamin et al.*** (***2016***). In the model used in this paper we assumed that all U1 individuals susceptible to RSV were at risk of their first RSV episode and that all O1 individuals had already been infected at least once, since re-infection within the same yearly epidemic is unlikely but nearly everyone has caught RSV by the age of two years old ***Glezen et al.***(***1986***).

### Conditional age of individuals

As mentioned above the high dimensionality of the RSV transmission model with two levels of social mixing was a limiting factor on the possible complexity of the compartmental framework representing the possible combinations of age and disease state. In order to both capture the structure of the population in households and incorporate finer-grained information about the ages of the modelled individuals, we used the conditional distributions to calculate the numbers of individuals in each finer-grained age category: (i) each month of first year of life, (ii) each year of life aged 1 - 18 and (iii) 18+ years old. We calculated empirical distributions for the number of individuals who were jointly in age category *a*, lived in a household of size *n*, which either contained at least one under one year old (*U* = 1) or not (*U* = 0), on days *t* = 1st Jan 2000, 2001,…, 2016, this joint distribution being denoted, ℙ_*t*_(*a, n, U*). We used these point estimates of the joint population distribution to represent the distribution for the year between point estimates. Conditional and marginal distributions for the finer-grained age category of an individual based on his crude age category *a <* 1 year or *a >* 1 year, his household size and whether the household contained an U1 or not were derived as standard, e.g.

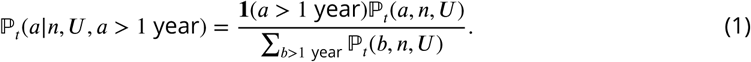

The conditional distribution for an individual’s household size and whether they lived in a household containing an U1 based on their age was constructed similarly. The reason we included a variable indicating whether the household of the individual contained an under one or not was because it was important to capture the pathway to transmission to the under-one year olds most at risk of disease due to contracting RSV. Because the exact birth dates where missing for a large number of people, and for model simplicity, we assumed that all U1 individuals aged to become O1 individuals at a constant rate 1 per year, which was equivalent to assuming that given that the exact age of an U1 individual was uniformly distributed between 0 and 1 years old, independently of the U1’s household configuration.

### Model Dynamics, forces of infection and susceptibility to RSV

The fundamental unit of the RSV transmission model developed for this paper was the household. Each household was described by the number of each type of individual inhabiting it, which we call the *household coniguration*. The type of individual within each household was identified by her RSV disease state and age category. The RSV transmission model described the dynamics of how many households were in each possible household configuration using an approach introduced by House and Keeling (***House and Keeling*** (***2008a***)). Mathematically, the number of households in a given household configuration at time *t* was denoted 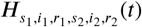, referring to the household configuration with exactly *s*_1_ U1 susceptibles, *i*_1_ U1 infecteds etc. In order to limit the number of possible household states we included only households of total size ten or less with two or less under ones. We chose these limits on the household size based on capturing ≈ 99% of the U1s in the population, and therefore the pathway to them catching RSV (*Supporting Information*). There were 1926 possible household configurations in the RSV transmission model. The vector ***H***(*t*) of number of households in each possible household configuration evolved according to the semi-linear ODE:

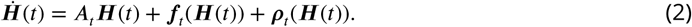

Each term describing the vector field of equation (2) corresponded to a dynamic component of the model:

1. RSV transmission within households, recovery of infected individuals, loss of immunity of recovered individuals, aging from U1 to O1 and turnover in household occupancy due to births and removals (*A*_*t*_***H***(*t*)).
2. RSV transmission between households due to age-group specific mixing (***f***_*t*_(***H***(*t*))).
3. Change in household numbers due to population flux, (***ρ***_*t*_(***H***(*t*))).

See *Supporting Information* for further details. The force of infection due to transmission within a household of generic configuration (*s*_1_, *i*_1_, *r*_1_, *s*_2_, *i*_2_, *r*_2_) was

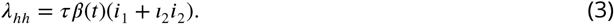

Where *τ* is the basic within-household transmission rate, *l*_2_ is the relative decrease in infectiousness of O1s compared to U1s, and *β* (*t*) is the seasonal variation in the transmission rate of RSV (see appendix 1). Transmission outside of the household within the wider community was assumed to be based on the finer-grained age categories introduced above. The conditional age distributions of the individuals allowed us to construct matrices (*P*_*H → A,t*_) to convert between the household configuration vector into a vector of number of infected individuals in each age category, weighted by their relative infectiousness, for any time *t* during the simulation: ***J*** (*t*) = *P*_*H →A,t*_***H***(*t*) (see *Supporting Information* for further details). The force of infection due to age-based mixing in the community was,

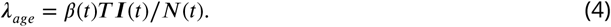

Where *T* was the community mixing rate matrix and *N*(*t*) was the total population size at time *t* ***Keeling and Rohani*** (***2008***). The force of infection on an individual within a given household was calculated using matrices constructed from the conditional distribution of an individual’s household type given their age, *λ*_*com*_ = *P*_*A → H,t*_*λ*_*age*_. The total force of infection on each individual was the sum of their infectious contact rates within the household and within the community, *λ* = *λ*_*hh*_ + *λ*_*com*_ + *λ*_*ext*_. Where *λ*_*ext*_ = *ϵβ* (*t*)/*N*(*t*) was the force of infection from outside KDHSS.

The actual infection rate for each individual was the force of infection ‘felt’ by the individual times the susceptibility of the individual. The susceptibility of under one year olds (*σ*_*U*1_) depended on whether or not the U1 individual was still protected from contracting RSV by maternally acquired antibodies, which we modelled as giving a random *M* days of protection; that is for an individual of age *A* days, *σ*_*U*1_ = 0 if *M > A* and *σ*_*U*1_ = 1 otherwise. Because RSV is strongly seasonal conditioning on an U1 not having received her first RSV infection does not strongly bias her age, therefore we used the mean susceptibility 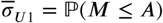 for under-ones. The susceptibility of over one year olds was chosen as if the individual had definitely received at least one RSV infection in the past, and definitely had no chance of being maternally protected (see *Supporting information*). We modelled the duration of maternal protection *M* as a truncated exponential distribution conditioned on being less than one year in duration; that is *M* ∼ exp(*α*) | (*M* ≤ 1 year).

### Hospitalisation rates

The chance of an infected individual becoming severely diseased after contracting RSV, and then seeking care at hospital, depended on that person’s age and number of infections ***Nokes et al.*** (***2008***); ***Ohuma et al.*** (***2012***). The conditional age of an U1 who has become infected was assumed to be independent of her household type (see above), but was shifted by conditioning on the age of the U1 being greater than her maternal protection period,

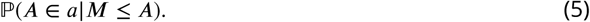

Which was calculated exactly (see *Supporting Information*). This took into account that increasing the duration of maternal protection would increase the age at infection and therefore reduce the risk of disease. O1s were assumed to have no maternal protection but their conditional age depended on their household type [equation (1)]. We used these conditional distributions to convert the incidence rate of U1s and O1s in each household type into dynamic incidence rates in each age category, ℐ_*a*_(*t*). By assuming that all O1s had been infected at least once we could use previously published age-dependent hospitalisation odds per infection *h*_*a*_ (***Kinyanjui et al.*** (***2015***) and *Supporting Information*) to determine the cumulative hospitalisations predicted by the model for each age category *a* and week interval *w*_*i*_ = (*t*_*i*,1_, *t*_*i*,2_),

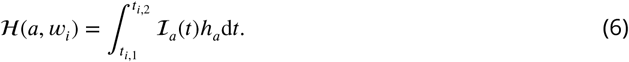

### Parameter Inference

The majority of the parameters for the RSV transmission model were drawn from the RSV literature (see *Supporting Information*) leaving four parameters, and the two hyperparameters of a normal distribution describing the random yearly variation in log-seasonality, to be inferred from hospitalisation data. The free parameters and distribution of the RSV transmission model were:

- Infectious contact rate outside the household between U1s and all others in the community accessing KCH (*b*_*U*1_).
- Infectious contact rate outside the household among all O1s in community (*b*_*O*1_).
- Infectious contact rate within the household (*τ*).
- Rate of loss of maternally derived immunity to RSV (*α*).
- The joint normal distribution of the yearly log-seasonality amplitude and phase ([*ξ, ϕ*] ∼ 𝒩 (***μ***, **Σ**)).

We also included an infectious contact rate for children of schooling age (5-18 years old; *b*_*S*_) which acted additionally to *b*_*O*1_; that is children of schooling age were at additional risk of contracting RSV on top of the risk due to mixing in the community. This meant that the mixing matrix in equation (4) was in block form,

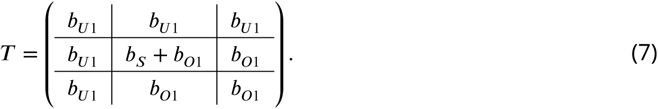

Where the blocks represented respectively under-one age categories, over-ones at school age categories and over-ones above school age categories. Unfortunately, we were unable to reliably identify *b*_*S*_ parameter jointly with the other parameters. Investigating a range of *b*_*S*_ values gave similar results for model fit and predictions for vaccine efficacy, the results in the main paper were for the highest value of *b*_*S*_ considered which was mildly pessimistic compared to *b*_*S*_ = 0 (see *Supporting Information*).

The data for parameter inference was RSV-confirmed, age-specific weekly admissions to Kilifi county hospital (KCH) hospitalisation data from September 2001 until September 2016 (see ***Nokes et al.***(***2009***) for study details). KCH serves as the primary care facility for the KHDSS population, and we assumed that all KHDSS members who accessed urgent hospital treatment due to RSV disease accessed their treatment at KCH. However, a significant number of admissions were from people not within the KHDSS survey leading to data re-scaling (see *Supporting Information*). The log-likelihood for a particular simulation corresponded to Poisson errors,

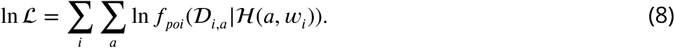

Where 𝒟 _*i,a*_ was the cumulative number of hospitalisation observed at KCH in age category *a* on week *w*_*i*_ and *f*_*poi*_(*x | μ*) is the probability mass function for a Poisson distribution with mean *μ*.

If the yearly realisations of the random seasonality (see appendix 1) were known, then the entire model would be deterministic and ln ℒ would be a function of the unknown parameters. Therefore, we treated the yearly variation in seasonality as missing data and used the Expectation-maximisation (EM) algorithm ***Dempster et al.*** (***1977***) to converge onto maximum likelihood estimates for the four free parameters, and the two hyperparameters of the log-seasonality model, 95% con1dence intervals were constructed using the likelihood profile technique (e.g. ***King et al.*** (***2008***) and *Supporting Information*).

### Modelling Vaccination

There were two vaccines used in this modelling study, which were deployed as part of the antenatal contact between pregnant women and skilled health professionals. We assumed that the maternal vaccine was delivered as one injection to the pregnant women in their third trimester. This achieved some unknown additional period of maternal protection, *P* days, on top of the random period *M*, that is after maternally vaccinating the period of protection became *M*_*vac*_ = *M* + *P*. Achieving an effective maternal vaccination coverage of *V*_*cov*_ shifted the mean susceptibility of U1s to 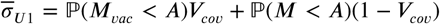, a linear increase in *V*_*cov*_. The change in distribution of age at infection was non-linear in *V*_*cov*_ because, conditional on an U1 being infected, it was more likely that the U1’s mother had not been vaccinated than the unconditional probability of non-vaccination, 1 - *V*_*cov*_ (see *Supporting Information*). We also assumed that there was a vaccine available that provoked an immune response in the vaccinated individuals similar to a natural infection; that is a susceptible *O*1 who is vaccinated immediately becomes ‘recovered’ and immune to RSV infection until her immunity waned. Immune response provoking vaccination was offered to all O1s in households when a birth occurred, as an addendum to the antenatal contact between mothers-to-be and health professionals. In principle, there were three dimensions to the coverage of the immunity provoking vaccine: (i) coverage of households, (ii) coverage within households, and (iii) vaccine effectiveness. For simplicity, we bundled these dimensions together, and vaccinated whole households at an effective vaccination rate (the product of the three dimensions of coverage).

### Model simulations

We simulated the model by numerically solving the high dimensional ODE [equation (2)] simultaneously with the ongoing cumulative hospitalisations in each age category, 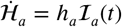, which allowed us to solve for the model predicted weekly hospitalisations [equation 6]. The initial state of the model was unknown. We initialised the model by starting with a completely susceptible population with the population demography set to mimic that of the KHDSS on 1st Jan 2000. We then simulated RSV transmission for 10 years, with demographic rates (e.g. birth rates) chosen to match those of KHDSS in year 2000 and the seasonal amplitude and phase of ln *β* set to their latest mean estimate, in order to equilibriate the state of the household model. Finally, we ran the model from 1st Jan 2000 until 1st September 2001. This provided the initial point for comparison to hospitalisation data. Numerical solutions were provided using the Sundials CVODE solver ***Cohen et al.***(***1996***) implemented within the **DifferentialEquations** package for Julia 0.6 ***Rackauckas and Nie*** (***2017***). For retrospective simulations comparing model predictions to data (Fig. 2) we used the most probable values of the yearly seasonality. For forecast simulations we generated 500 realisations of yearly seasonality over 10 years from the distribution inferred in model inference, this gave 500 predictions for the time series of future hospitalisations. We typically presented medians of these predictions (e.g. Fig. 4).

## Acknowledgments

This work was funded by the Wellcome Trust (Grant ref 102975), and was published with permission of the Director of KEMRI. Kat Rock kindly supplied some clipart for plotting.

## Appendix 1

**Modelling seasonality in RSV transmission among KDHSS**

RSV is a seasonal virus, in temperate climates the peak month for RSV incidence tends to be consistent year-on-year. Therefore, modelling approaches aimed at understanding RSV transmission in temperate climates have used an annually periodic deterministic function, with the timing of peak infectiousness of RSV being either a model parameter ***Yamin et al.*** (***2016***) or itself a function of climatic variable to be fitted using regression methods ***Pitzer et al.***(***2015***).

The seasonal drivers of RSV transmission in the tropics are less clear ***Paynter*** (***2015***). At KCH the most common trough month for RSV hospitalisations was September, which lead us to define the RSV ‘year’ as September - September. The most common month for peak hospitalisation in each RSV year was January, however there was significant variation in peak month between RSV seasons with peaks occurring in each month November - April between 2002-2016 (Fig 1).

**Appendix 1 Figure 1.**
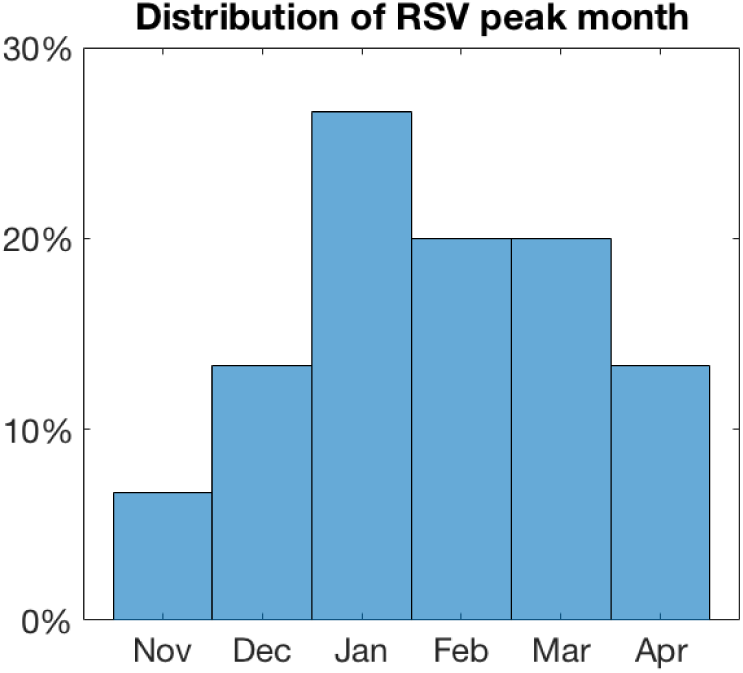
Distribution of peak month for RSV hospitalisations at KCH.

The year-on-year variation in peak month for RSV hospitalisation means that naively inferring a single fixed peak infectiousness parameter would not be a successful inference strategy. However, determining the precise mechanistic reason for shifting seasonality was challenging for the KDHSS population. RSV has been positively associated with the rainy season in some tropical settings ***Paynter et al.*** (***2012***); ***Paynter*** (***2015***), however this is not obviously the case in Kilifi county where the rainy season is April to June with short rains October to December. There have been many proposed mechanisms for erratic periodicity in transmission (for a wide variety of infectious pathogens) which *could* be relevant to RSV transmission in Kilifi, for example, dynamical attractor switching ***Keeling et al.*** (***2001***), or the effect of species/strain interaction ***Bhattacharyya et al.*** (***2018***). In particular, strain competition between RSV A and RSV B has been identified a mechanism for generated complex seasonal dynamics ***White et al.***(***1999***).

In this paper, we took an agnostic view and rather than choosing a mechanistic hypothesis for erratic seasonality from the many possible, we assume that the time-varying infectiousness of RSV alters randomly (but from a common distribution) year to year:

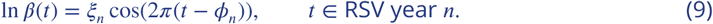

Where the RSV infectiousness (*ξ*_*n*_) and seasonal peak timing (*ϕ*_*n*_) for each RSV year *n* are drawn jointly from a normal distribution common to each year (*ξ* _*n*_, *ϕ* _*n*_) ∼ 𝒩 (***μ***, **Σ**). During model inference the yearly *ξ*_*n*_ and *ϕ*_*n*_ realisations are treated as latent variables; their mean and covariance matrix are imputed along with other model parameters.

