## Supplementary information for "Reducing RSV hospitalisation in a lower-income country by vaccinating mothers-to-be and their households"

### Household- and age-structured RSV transmission model details

As described briefly in the main text, we developed a dynamic model for simulating the spread of RSV through the KDHSS population. The model was a hybrid between a mechanistic ODE approach, this included detailed household structure but only a simplified set of age-and-disease states for individuals within the households, and a data-driven empirical model, this used the observed joint distributions of KDHSS individuals' household occupancy and ages to generate conditional predications of individual detail beyond that of the mechanistic part of the model.

#### Brief comparison to age-structured RSV transmission models

A commonly used conceptual framework for modelling epidemic transmission with a population is the compartmental model [1, 2]; each person's disease state is described as being one of a finite number of possibilities, e.g. susceptible, infectious, recovered, which define that person's risk of contracting the infectious pathogen or transmissibility whilst infected with the pathogen. Additionally, it is usually important to capture the heterogeneity of the population, also called the *population structure*[1], and therefore each person will be described by their position in the population with sufficient detail that a rate of contact can be modelled between any pairs of individuals, see Diekmann and Heesterbeek for a more detailed discussion on modelling population structure [3]. RSV transmission models have most commonly used age structure to describe heterogeneity in the population; each individual is described jointly by their disease state and which age interval (from some predetermined set of intervals) they occupy [4, 5, 6]. For age-structured RSV transmission models there are two dynamical elements: the transmission of disease and the demographic turnover of the population (births, deaths and ageing). At the level of the individual these are modelled as discrete random events occurring at some per-capita rate [7]. However, for large populations, there will be a very large number of individuals in each age-and-disease state, and the flux of population density in each age-and-disease state converges in probability onto the solution of a set of ordinary differential equations (ODEs) as the population size is treated as converging to infinite size [8, 9, 3]. The limiting ODE model has as many degrees of freedom as there are age-and-disease state combinations in the epidemic model. In most epidemic modelling studies it is the deterministic evolution of the solution to these ODEs that is usually given as the transmission

[1] In contrast to unstructured populations where every individual is treated as interchangeable.

model description.

In this paper, the essential modelling concept was to shift the focus away from numbers of individuals in each age-and-disease state and towards the number of households in each possible *household configuration*. A household configuration describes the number of individuals in each age-and-disease state who cohabit within a single household. Including households within the model adds a potentially relevant layer of realism; the social contacts within a household are *persistent*, therefore pairs of individuals that cohabit will repeatedly have the opportunity to infect one another if RSV enters the household but be relatively cocooned from infection if RSV has not entered the household. Age-structured transmission models implicitly assume that no two individuals contact one another more than once. To see this consider a population size of $N$; the rate of any individual contacting another single individual is $\mathcal{O}(1/N)$ therefore the probability that an individual selects the same other individual twice for contact over any finite time horizon goes to zero as $N \to \infty$ (which is also the limit at which the ODE model is valid). For household models the discrete random events that change the state of individuals (infection, death etc.) also change the household configuration. When the number of households is very large, there will be a large number of households in each possible household configuration and, as with age-structured models, there is convergence onto a set of ODEs with as many degrees of freedom as the number of possible household configurations.

The possible household configurations, or *state space*, of a household- and age-structured RSV transmission model is considerably larger than it would be for the equivalent age-structured model. If there are $m$ possible age-and-disease states then the number of possible household configurations for a household of size $n$ is given by a standard combinatorial identity, $\binom{n+m-1}{n}$. In thus paper we consider a range of household sizes up to a maximum size $n_{max}$, therefore the number of household configurations was,

$$\text{\# household configurations} = \sum_{n=1}^{n_{max}} \binom{n+m-1}{n}.$$

The number of possible household configurations grows very rapidly [Fig. 1]. Therefore, having a sufficiently large $n_{max}$ to capture the target population required using a relatively simple compartmental age-and-disease state model for RSV infection.

#### Age-and-disease states for the household model

A literature review of mechanistic RSV transmission models revealed a number of critical common features:

- At birth newborns are born protected against RSV infection due to antibodies gained from their mother via trans-placental transfer. This is typically modelled as a maternally protected disease state $M$ e.g. [6].
- The probability of developing severe disease and being hospitalised depends on a person's age, and number of times infected in the past, e.g. [5].
- The susceptibility to RSV infection per infectious contact, their infectiousness after infection, and the expected time taken to become recovered from RSV depend on number of times previously infected, e.g. [5].

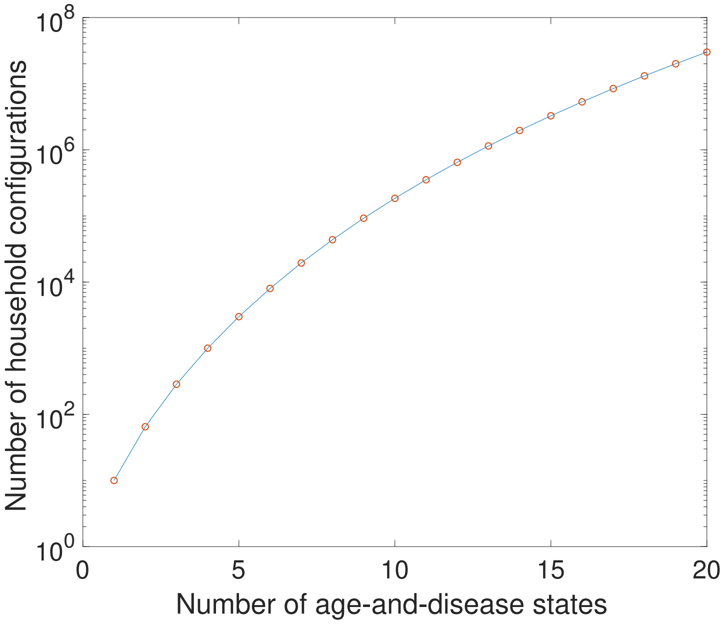


Figure 1: Growth in number of possible household configurations as complexity of the underlying age-and-disease state model grows. Calculated for a maximum household size of 10.

The high dimensionality of household- and age-structured models necessitated using the most minimal age-and-disease state model possible for RSV (see above). To do this we use an extremely parsimonious approach. The possible age-and-disease state for individuals are: susceptible *or* maternally protected and under the age of one ($S_1$), infectious and under the age of one ($I_1$), recovered and under the age of one ($R_1$), susceptible and over the age of one ($S_2$), infectious and over the age of one ($I_2$) and recovered and over the age of one ($R_1$). An under-one year old (U1) experiencing some force of infection $\lambda$ becomes infected ($S_1 \to I_1$) and infectious to RSV at a rate $\sigma_{U1}\lambda$ where $\sigma_{U1}$ is the average susceptibility of an U1 year old to RSV. After becoming infected the U1 ceases to become infectious at a rate $\gamma_1$ ($I_1 \to R_1$) and then is immune to reinfection to RSV for a period of time. The immunity derived from natural infection is lost at a rate $\nu$, and the U1 revert to susceptibility but in the $S_2$ category ($R_1 \to S_2$). The reason we transition recovered U1s to a susceptible over-one year old (O1) is that due to the seasonality of RSV it is very rare for a person to be infected more than once in one epidemic season, therefore functionally by the time an individual is facing the risk of their second RSV lifetime infection they will very likely be over one. All U1s age at the rate $\eta = 1/365.25$ days$^{-1}$ becoming individuals in the same disease state but over-one ($S_1 \to S_2$, $I_1 \to I_2$, $R_1 \to R_2$). An O1 individual experiencing a force of infection $\lambda$ becomes infected and infectious ($S_2 \to I_2$) with RSV at a rate $\sigma_{O1}\lambda$ where $\sigma_{O1}$ is the relative susceptibility of O1s compared to an U1 no longer protected by maternal antibodies. Infectious O1s cease being infectious ($I_2 \to R_2$) at a faster rate than U1s, $\gamma_2 > \gamma_1$, but revert to susceptibility ($R_2 \to S_2$) at the same rate $\nu$ [Fig. 2].

As mentioned in the main document we relate this simple age-and-disease state model to more complicated RSV models by (i) using the conditional age distribution of individuals to address questions that required a more complicated age structure than a simple under/over-one binary choice, for example whether susceptible under ones were still protected by maternal antibodies, and (ii) by assuming that all over-ones have been infected at least once and all susceptible U1s have never been infected and *might* still be protected by maternal antibodies.

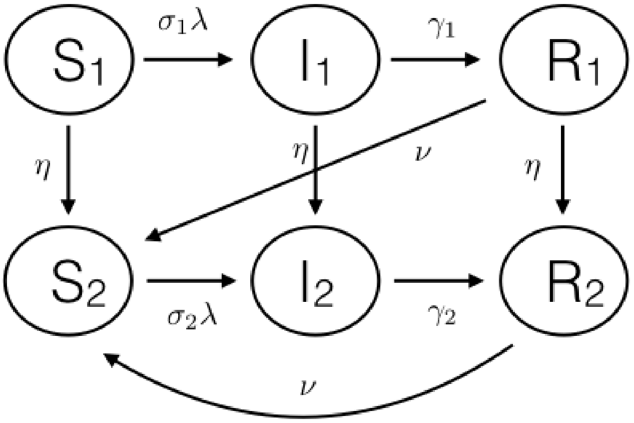


Figure 2: Schematic diagram of the basic age-and-disease state compartmental model for the individuals inside the households.

#### Conditional age of individuals

The household-structured RSV transmission model has a crude age category approach (U1/O1); however, we recover much more detail about the population by using conditional age distributions. The point of including extra age categories beyond the simple U1/O1 divide was that as well as modelling household based processes we can also model age-based processes like mixing outside the household, the chance of an U1 having finished her maternal protection period, and chance of disease if infected.

As mentioned in the main text we inferred an empirical joint distribution for individuals of their household size $n$, a boolean indicating whether that household contained an U1 member or not (always true if the individual is U1) and their age category $a$. The age categories were chosen so as be interpretable in the context of other published work on RSV disease risk, in particular Kinyanjui *et al* [5]. The age categories were: (i) each month of first year of life, (ii) each year of life aged 1 - 18 and (iii) 18+ years old. The exact birth date was often unavailable and given as the first day of birth. Therefore, for simplicity, and in line with maximum entropy considerations, we assumed that all under-one year olds had a birth date drawn uniformly across their year of birth.

For any given day $t$ this joint distribution was calculated empirically by counting the number of individuals in all three categories relative to the total number of individuals alive,

$$\mathbb{P}_t(a, n, U) = \frac{\#\text{individuals in KDHSS on day } t \text{ in age category } a\text{, living in house size } n\text{, with U1 member boolean } U}{\#\text{individuals in KDHSS on day } t}. \quad (1)$$

Since these joint distributions varied over time we calculated empirical distributions on days $t$ = 1st Jan 2000, 2001,..., 2016, and used these as representative for that year. Calculating the joint distributions allowed calculation of any conditional distributions, e.g.

$$\mathbb{P}_t(a|n,U,a<1\text{ year}) = \frac{\mathbf{1}(a<1\text{ year})}{T}\int_0^T \mathbf{1}(t\in a)\,\mathrm{d}t, \quad (2)$$

$$\mathbb{P}_t(a|n,U,a>1\text{ year}) = \frac{\mathbf{1}(a>1\text{ year})\mathbb{P}_t(a,n,U)}{\sum_{b>1\text{ year}}\mathbb{P}_t(b,n,U)}, \quad (3)$$

$$\mathbb{P}_t(n,U|a) = \frac{\mathbb{P}_t(a,n,U)}{\sum_b \mathbb{P}_t(b,n,U)}. \quad (4)$$

Where $T$ is the duration of one year in the model units (we used days) and "$a<1$ year" means that the age category $a$ is completely contained in the first year of life.

#### Household- and age-structured model dynamics

A *household configuration* is a tuple of the number of individuals in each age-and-disease state who cohabit a household. The generic household configuration is denoted $h=(s_1,i_1,r_1,s_2,i_2,r_2)$, indicating that the household has precisely $s_1$ individuals in state $S_1$, $i_1$ individuals in state $I_1$ etc. The *household* size is the number of people living in the household (i.e. $s_1+i_1+r_1+s_2+i_2+r_2$). We denote the space of possible household configurations $\Sigma$ and number of households in the state $h$ at time $t$ as $H_h(t)$. It is useful to consider a vector quantity over all possible household configurations such as $\boldsymbol{H}(t)=(H_h(t)\mid h\in\Sigma)$ where we have generated some ordering for elements $h\in\Sigma$. It is clear that the knowledge of $(\boldsymbol{H}(t),t\geq 0)$ would allow us to reconstruct the dynamics of individuals. For example, using the function $f(h)=s_1$ for each $h\in\Sigma$ in a vectorised form $\boldsymbol{f}=(f(h)\mid h\in\Sigma)$ allows us to track the dynamics of numbers of $S_1$ individuals: $(\boldsymbol{f}\cdot\boldsymbol{H}(t),t\geq 0)$.

As mentioned above, age-structured models are constructed by considering the per capita rate of events affecting the state of individuals. Household- and age-structured models are constructed by considering the per household rate of events that affect the household configuration (see [10] for further mathematical details). In the following we list the events that change the household model divided into three groups: events due to transmission within the household, events due to transmission between households and events due to demographic turnover.

##### Seasonality in transmission

As described in the appendix to the main text, RSV transmission in Kenya peaks annually, most of the cases recorded between 2002-2016 at KDH were observed during January, and the least during September (figure 3). However, the timing of the peak each year has shown significant variation over that time, ranging from November to April as peak months. To account for both the annual peak in RSV incidence, and also the unknown drivers of seasonal peak timing, we allowed each year of RSV transmission (we define the RSV season as beginning on September 1st) to have a random amplitude $\xi_i$ and phase $\phi_i$ where $i$ indexes the seasons. These random seasonal variables are not observed directly, so we assume that each pair is jointly normal $(\xi_i,\phi_i)\sim\mathcal{N}(\boldsymbol{m},\Sigma_{\xi\phi})$ where the mean vector $\boldsymbol{m}=[m_\xi\ m_\phi]$ and covariance matrix $\Sigma_{\xi\phi}$ will be inferred jointly with other model parameters. For each RSV season we scale infection rates by the time varying seasonality factor:

$$\ln\beta(t)=\xi_i\cos(2\pi(t-\phi_i)/365.25),\qquad \text{for } t \text{ in season } i. \quad (5)$$

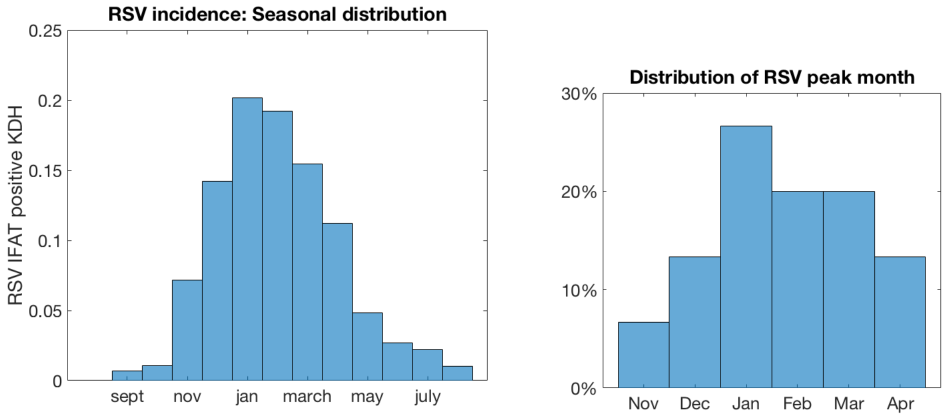


Figure 3: Distribution of total incidence by month of year (*left*) and peak incidence (*right*).

##### Events due to RSV transmission within the household

- Infection of susceptibles from within the household:

$$\text{For U1s: } [s_1, i_1, r_1, s_2, i_2, r_2] \to [s_1 - 1, i_1 + 1, r_1, s_2, i_2, r_2] \text{ at rate: } \sigma_{U1}\beta(t)\tau s_1(i_1 + \iota_2 i_2), \quad (6)$$
$$\text{For O1s: } [s_1, i_1, r_1, s_2, i_2, r_2] \to [s_1, i_1, r_1, s_2 - 1, i_2 + 1, r_2] \text{ at rate: } \sigma_{O1}\beta(t)\tau s_2(i_1 + \iota_2 i_2). \quad (7)$$

$\tau$ is the household infection rate, $\iota_2$ is the reduction in infectiousness due to being an O1, $\beta(t)$ is the seasonally varying component to the transmission rate and $\sigma_{O1}$ is the reduction in susceptibility due to being O1. Note that the true infection rate for U1s is $\sigma_{U1}\lambda_{hh}$ and for O1s is $\sigma_{O1}\lambda_{hh}$ as defined in main text. $\sigma_{U1}$ is the probability that an U1 individual is no longer protected by maternal antibodies, calculated by integrating over the individuals conditional age distribution as follows. Maternal protection was assumed to be 100% effective but only for a random duration per newborn of $M$ days, therefore using the uniform age distribution conditional on the individual being under one years old (see above),

$$\sigma_{U1} = \frac{1}{T}\int_0^T \mathbb{P}(M \leq a)\,\mathrm{d}a. \quad (8)$$

Where $T$ is the duration of a year expressed in the units of the simulation (we used days so $T = 365.25$ days). The probabilistic model for the duration of maternal protection was $P \sim \exp(\alpha)|M \leq T$ days , where $\alpha$ is the waning maternal immunity rate. The distribution function for $M$ is

$$\mathbb{P}(M \leq a) = \begin{cases} (1 - \exp(-a/\bar{M}))/(1 - \exp(-T/\bar{M})) & 0 \leq a \leq T \\ 1 & \text{otherwise} \end{cases} \quad (9)$$

Where $\bar{M} = 1/\alpha$ is the mean period of maternal protection without conditioning on $M \leq T$, the true mean period of protection is $\mathbb{E}[M] = \bar{M} - T/(e^{T/\bar{M}} - 1)$ but this turns out to be a very small correction to $\overline{M}$ since we fit to $\overline{M}$ being less than 30 days (see below), therefore for simplicity we call $\overline{M}$ the mean duration of maternal protection to RSV. Substituting into equation (8) and direct integration gives,

$$\sigma_{U1} = \frac{1}{1 - e^{-T/\bar{M}}} - \frac{\bar{M}}{T}. \quad (10)$$

Note that $\sigma_{U1} \approx 1 - \bar{M}/T$ when $\bar{M} \ll T$.

- Recovery of infecteds:

$$\text{For U1s: } [s_1, i_1, r_1, s_2, i_2, r_2] \to [s_1, i_1 - 1, r_1 + 1, s_2, i_2, r_2] \text{ at rate: } \gamma_1 i_1, \quad (11)$$
$$\text{For O1s: } [s_1, i_1, r_1, s_2, i_2, r_2] \to [s_1, i_1, r_1, s_2, i_2 - 1, r_2 + 1] \text{ at rate: } \gamma_2 i_2. \quad (12)$$

Where $\gamma_1$ and $\gamma_2$ are the recovery rates of U1s and O1s.

- Reversion to susceptibility:

$$\text{For U1s: } [s_1, i_1, r_1, s_2, i_2, r_2] \to [s_1, i_1, r_1 - 1, s_2 + 1, i_2, r_2] \text{ at rate: } \nu r_1, \quad (13)$$
$$\text{For O1s: } [s_1, i_1, r_1, s_2, i_2, r_2] \to [s_1, i_1, r_1, s_2 + 1, i_2, r_2 - 1] \text{ at rate: } \nu r_2. \quad (14)$$

Where $\nu$ is the reversion to susceptibility/waning immunity rate.

##### Events due to RSV transmission from without the household

In a purely age-structured transmission model the number of RSV infecteds in each age category, $\boldsymbol{I}(t) = (I_a(t))_{a\in\mathcal{A}}$, is a dynamic model variable which evolves according to a set of ODEs. For the household- and age-structured model we derived $\boldsymbol{I}(t)$ from the household configuration dynamics and the conditional age distributions as the expected number of infecteds in each category given the distribution of household configurations $\boldsymbol{H}(t)$. Note that knowing a household configuration specifies both the household size $n = s_1 + i_1 + r_1 + s_2 + i_2 + r_2$ and the under-one occupant boolean $U = \mathbf{1}(s_1 + i_1 + r_1 > 0)$. Therefore, we could define a $|\mathcal{A}| \times |\Sigma|$ conversion matrix to convert between the dynamic $\boldsymbol{H}(t)$ variables into the implied $\boldsymbol{I}(t)$ variables,

$$P_{H\to A,t} = (\mathbb{P}_t(a|h))_{a\in\mathcal{A}, h\in\Sigma}, \quad (15)$$
$$\boldsymbol{I}(t) = P_{H\to A,t}\boldsymbol{H}(t). \quad (16)$$

The age dependent force of infection on each individual in age category $a$, $\lambda_{age}(a)$ depends on a community age mixing matrix $T = (T(a,b))_{a\in\mathcal{A}, b\in\mathcal{A}}$,

$$\lambda_{age}(a,t) = \sum_{b\in\mathcal{A}} T(a,b)[\mathbf{1}(a < 1 \text{ year}) + \iota_2 \mathbf{1}(a > 1 \text{ year})] I_b(t)/N(t). \quad (17)$$

Where $N(t)$ is the total population size at time $t$. This is a standard formulation for force of infection between different age groups (see Keeling and Rohani [2]). In principle any age-mixing matrix can be used as $T$, however we use a simple matrix in block form that differentiated only between U1s, O1s of school age, and all other O1s (see main text). The force of infection on U1 and O1 individuals within households was calculated using a $|\Sigma| \times |\mathcal{A}|$ conversion matrix, and a small force of infection from outside the KDHSS was added, $\epsilon$,

$$P_{A\to H,t} = (\mathbb{P}_t(h|a))_{h\in\Sigma, a\in\mathcal{A}}, \quad (18)$$
$$\lambda_{com}(U1,h,t) = \sum_{a<1 \text{ year}} \mathbb{P}_t(h|a)\lambda_{age}(a,t) + \epsilon/N(t), \quad (19)$$
$$\lambda_{com}(O1,h,t) = \sum_{a>1 \text{ year}} \mathbb{P}_t(h|a)\lambda_{age}(a,t) + \epsilon/N(t). \quad (20)$$

The external infection event changes the household configuration:

- Infection of susceptibles from outside the household:

$$\text{For U1s: } [s_1, i_1, r_1, s_2, i_2, r_2] \to [s_1 - 1, i_1 + 1, r_1, s_2, i_2, r_2] \text{ at rate: } \sigma_{U1}\beta(t)s_1\lambda_{com}(U1,h,t), \quad (21)$$
$$\text{For O1s: } [s_1, i_1, r_1, s_2, i_2, r_2] \to [s_1, i_1, r_1, s_2 - 1, i_2 + 1, r_2] \text{ at rate: } \sigma_{O1}\beta(t)s_2\lambda_{com}(O1,h,t). \quad (22)$$

#### Events due to demographic change in the population

In the household-and-age-structured RSV model we track demographic change both by using the yearly updated joint distributions of age and household size and by the dynamics of the household configurations $\boldsymbol{H}(t)$. The number of households of each size $n$ changed over time due to the effect of people leaving home, births, deaths, out-migration from KDHSS and in-migration into KDHSS. Moreover, the mean number of U1s per household of each size evolved over time. Rather than track all the possible events that change the demography of the KDHSS, we focus on (i) the ageing of the U1s becoming O1s, (ii) capturing the household size dependent birth rate, and (iii) capturing the change in household numbers for each household size.

The recorded birth rate that can be inferred from the KDHSS data set included newborns who out-migrate, neglected newborns that in-migrate at a very young age, and obviously some newborns die whilst very young. As mentioned above, we did not mechanistically track every possible demographic event, but instead calculated the *effective* birth rate that arrived at the correct mean number of U1s for each household size. For simplicity, we assumed that the effective birth rate was a *turnover rate* for households; that is each birth is associated with a per-capita rate of an O1 leaving the household. This arrived at the correct density of U1s in the population, and in each size group of households, at the cost of assuming that events occurred at the same time rather than at the same rate.

The number of households of each size changed over time as the overall population size changed and individuals left households in order to form new households. As with the demographic turnover rate, there were multiple different mechanisms whereby new individuals entered the population and formed new houses or individuals and groups left the population, e.g. whole groups arrived and formed a new house, individuals arrived and joined individual houses etc. Moreover, the RSV infection status of the new entrants to the population were unknown. We assumed that new entrants arrived as households with the same distribution of household configurations as already observed in the population; that is that new arrivals didn't have a net effect on the *proportion* of individuals in each age-and-disease state just by arriving, although obviously as the population grew this has an effect of the number of hospitalisations we expected.

The demographic events that changed the household configurations were:

- Aging:

$$[s_1, i_1, r_1, s_2, i_2, r_2] \to [s_1 - 1, i_1, r_1, s_2 + 1, i_2, r_2] \text{ at rate: } \eta s_1, \quad (23)$$
$$[s_1, i_1, r_1, s_2, i_2, r_2] \to [s_1, i_1 - 1, r_1, s_2, i_2 + 1, r_2] \text{ at rate: } \eta i_1, \quad (24)$$
$$[s_1, i_1, r_1, s_2, i_2, r_2] \to [s_1, i_1, r_1 - 1, s_2, i_2, r_2 + 1] \text{ at rate: } \eta r_1. \quad (25)$$

  Where $\eta = 1/T$ is the aging rate at which U1s become O1s. $T$ is the duration of a year expressed in the units of the simulation (we used days so $T = 365.25$ days).

- Demographic turnover due to births and O1s leaving their household:

$$[s_1, i_1, r_1, s_2, i_2, r_2] \to [s_1 + 1, i_1, r_1, s_2 - 1, i_2, r_2] \text{ at rate: } \mu(n,t) s_2, \quad (26)$$
$$[s_1, i_1, r_1, s_2, i_2, r_2] \to [s_1 + 1, i_1, r_1, s_2, i_2 - 1, r_2] \text{ at rate: } \mu(n,t) i_2, \quad (27)$$
$$[s_1, i_1, r_1, s_2, i_2, r_2] \to [s_1 + 1, i_1, r_1, s_2, i_2, r_2 - 1] \text{ at rate: } \mu(n,t) r_2. \quad (28)$$

  If there is at least one O1 left in the household, the birth/turnover rate is zero for households with only 1 O1; that is there are never any households of only U1s. $\mu(n,t)$ is the turnover rate per O1 household member in a household of size $n$ at time $t$ replacing them with susceptible U1s for households of size $n$. The turnover rates for each year were chosen so that the correct density of

U1s per household was achieved (approximately). Following is a description of the fitting process so that the turnover rate lead to this household demography:

1. *Collect the empirical distribution of U1s per household size.* For each household size $n = 1, \dots, n_{max}$ we calculated the mean number of U1s per household at $y =$ 1st jan 2000-2017, this was denoted: $\overline{N}_{U1}(n, y)$.
2. *Calculate the implied distribution of U1s per household size for any given birth/turnover rate.* For any given birth/turnover rate, $\mu$, the equilibrium probability of finding $k$ U1s in a household of size $n$ is

$$\pi(k|n,\mu) \propto \left(\frac{\mu}{\eta}\right)^k \binom{n}{k} \qquad k = 0, ..., n-1, \tag{29}$$

$$\pi(n|n,\mu) = 0. \tag{30}$$

   Equation (29) is just the equilibrium distribution of a birth-death process [11].
3. *Matching the empirical distribution to the implied distribution.* We used a root-finder to find the turnover rate that matches the simulation's mean number of U1s per household of each size to the empirical data, *for the next year*:

$$\mu(n,t) \text{ is the solution to } \sum_{k=0}^{n-1} k\pi(k|n,\mu(n,t)) = \overline{N}_{U1}(n, y+1) \text{ for all } t \text{ in year } y. \tag{31}$$

- Change in number of households due to population flux

$$[s_1, i_1, r_1, s_2, i_2, r_2] \to 2[s_1, i_1, r_1, s_2, i_2, r_2] \text{ at rate: } \frac{r(n,t)}{\sum_{h\in\Sigma_n} H_h(t)}, \text{ if } r(n,t) \geq 0, \tag{32}$$

$$[s_1, i_1, r_1, s_2, i_2, r_2] \to \emptyset \text{ at rate: } \frac{|r(n,t)|}{\sum_{h\in\Sigma_n} H_h(t)}, \text{ if } r(n,t) < 0. \tag{33}$$

(34)

Where, $\Sigma_n = \{h = [s_1, i_1, r_1, s_2, i_2, r_2] \mid s_1 + i_1 + r_1 + s_2 + i_2 + r_2 = n\}$ was the set of household configurations of households of size $n$. $r(n,t)$ was the daily rate of change of number of households of size $n$ interpolated between the empirical distribution dates.

#### Simulating the model

The model above could in principle have an infinite number of states if the household size was not limited (see above). We chose limits on the household size based on capturing $\approx$ 99% of the U1s in the population, and therefore the pathway to them catching RSV. The limits were: (i) no household is bigger than size 10, and (ii) no household has more than 2 U1s. This also covers the big majority of the total numbers of households (see figure 4). The $n_{max} = 10$ limit was imposed by initialising the model without households of size $> 10$, and setting $r(n,t) = 0$ for all $n > 10$. The $\leq$ 2 U1 limit was imposed by setting the birth/turnover rate to zero for all households with 2 U1s. Putting the limits in reduces the dimensionality of the system to 1926 different household configurations.

Note that the events that either change a household's configuration or change the number of households described above can be divided into two categories: [1] those with rates that only depended on the household's configuration, e.g. infection within the household, or ageing of U1s, and, [2] those with rates that depended on the configurations of other households, e.g. transmission between households

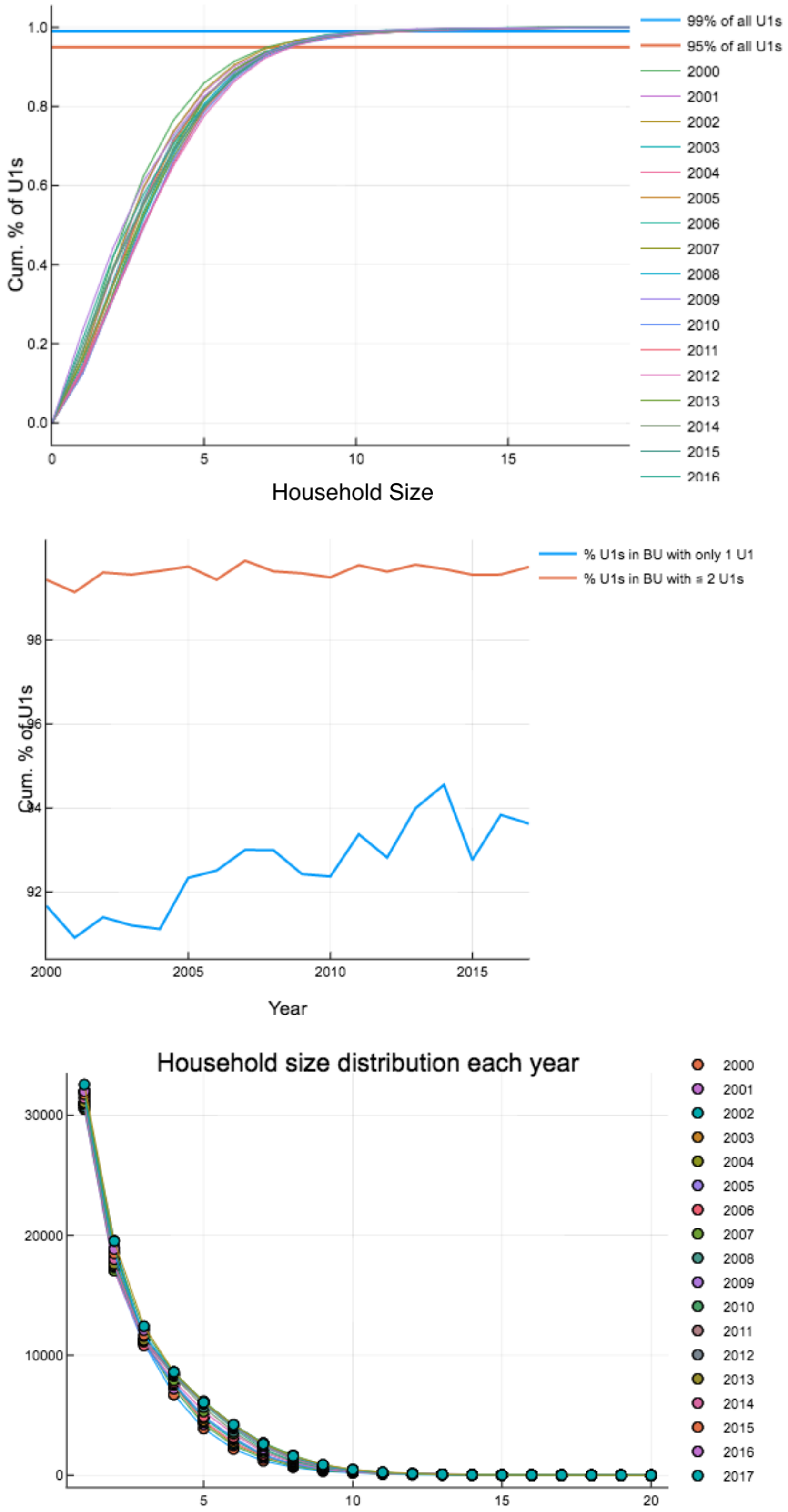


Figure 4: Household occupancy characteristics calculated on each 1st Jan 2000-2017. *Top:* Percentage of U1s in households of a certain size or smaller. *Middle:* Percentage of U1s in households with only one U1 and households with one or two U1s. *Bottom:* Household size distribution.

or the rate of change of household numbers. The events in category [1] translate to linear dynamics for $\boldsymbol{H}(t)$, events in category [2] translate to non-linear dynamics [12]. Overall, the dynamics of $\boldsymbol{H}(t)$ obey the semi-linear dynamical system,

$$\dot{\boldsymbol{H}}(t) = A_t \boldsymbol{H}(t) + \boldsymbol{f}_t(\boldsymbol{H}(t)) + \boldsymbol{\rho}_t(\boldsymbol{H}(t)). \tag{35}$$

$A_t$ is a matrix which encodes the dynamics of events in category [1], $\boldsymbol{f}_t(\boldsymbol{H}(t))$ encodes the transmission between households, and $\boldsymbol{\rho}_t(\boldsymbol{H}(t))$ encodes the rate of change of numbers of households in each configuration. We initialised the dynamics of equation (35) by starting with a completely susceptible population, allowing RSV to be introduced via the external force of infection and running for 10 years (see main text).

Equation (35) has two properties that are important to note:

- The change rate in households of size $n$ is independent of the transmission dynamics:

$$\partial_t \Big( \sum_{h \in \Sigma_n} H_h(t) \Big) = r(n,t), \qquad n = 1, \dots, 10. \tag{36}$$

- The dynamics of the proportion of households in a given state $P_h(t) = H_h(t)/\sum_{h'} H_{h'}(t)$ is not directly affected by the change rates ($\boldsymbol{\rho}_t$) in households:

$$\partial_t \boldsymbol{P}_t = A_t \boldsymbol{P}_t + \frac{\boldsymbol{f}_t\Big(\boldsymbol{H}_t\Big)}{\sum_{h'} H_{h'}(t)} \tag{37}$$

Equations (36) and (37) guarantee the desired modelling features discussed above. Equation (36) gives that the change in the number of households of each size matches the empirical rate of change for each year, we also verified this by numerical solution of equation (35) (figure 5). Equation (37) shows that the rate of change of household numbers doesn't directly effect the proportion of households in any given configuration. We also verified that the number of U1s and O1s was close to their empirical values (figure 6).

Equation (35) was difficult to solve efficiently because it is both numerically stiff and high dimensional. We numerically solved equation (35) using the Julia **DifferentialEquations** package implementation of the CVODE solver, with an efficient Krylov method (GMRES) to solve the implicit timestepping (see main text). We also used the **DifferentialEquations** efficient event handling which allowed us to change parameters (like the household change rate) at specific times without damaging the performance of the solver, or having to restart simulations.

#### Parameters for the household- and age-structured RSV transmission model

The parameters for the household- and age-structured transmission model were drawn from four sources:

- A literature review of infectiousness duration and other epidemiological quantities; table 1.
- Calculated from the empirical joint distributions (see above), i.e. equation (1), or inherent to the dynamics; table 1.
- Age-dependent hospitalisation probability per RSV infection derived from Kinyanjui *et al* [5]; table 2. Hospitalisation probability was the probability that an infected individual would develop severe

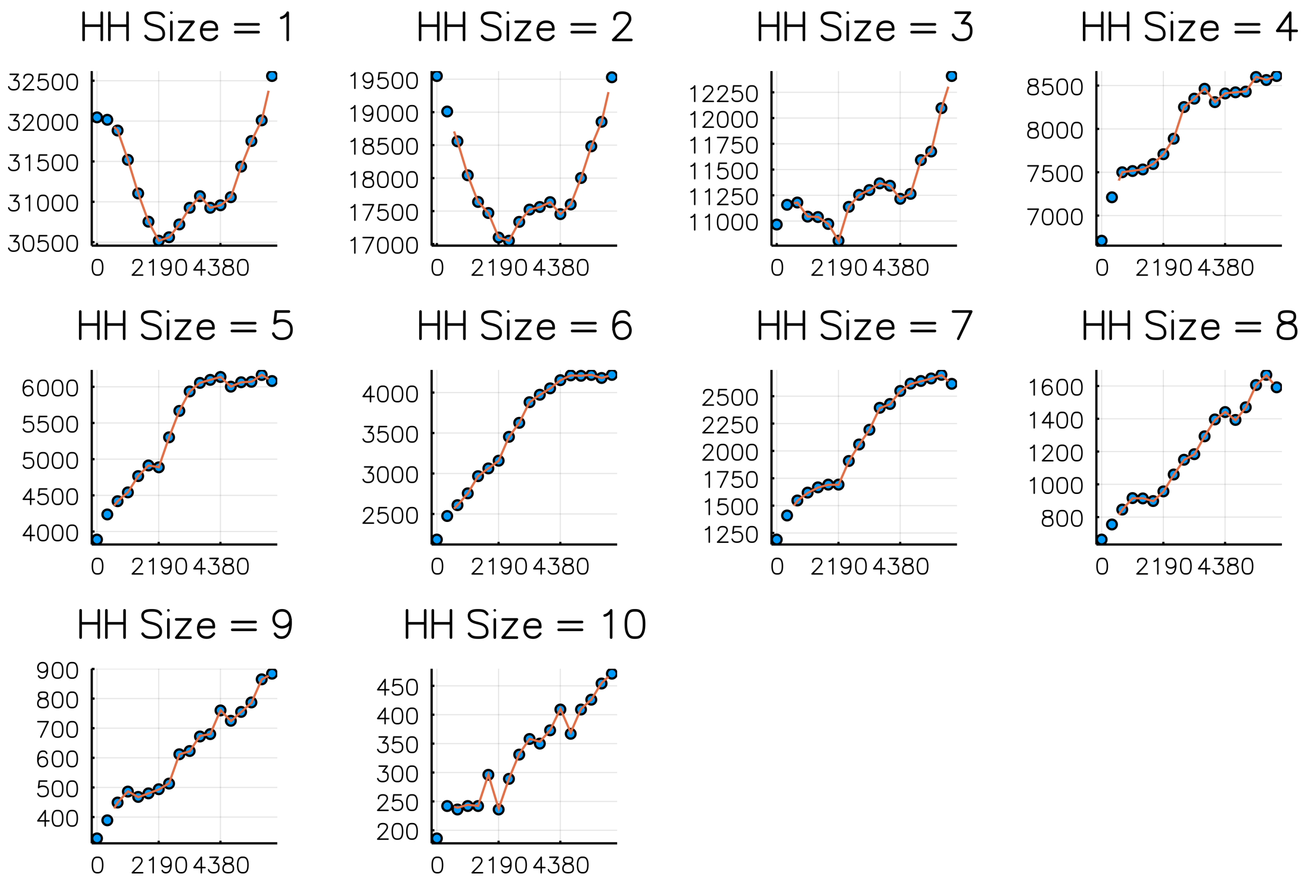


Figure 5: Comparison of numbers of households of sizes 1-10 on each 1st Jan 2000-2017 (dots) against simulated values (curve). Simulation is from Sept 2001 - Sept 2016. Horizontal axis is days since 1st Jan 2000.

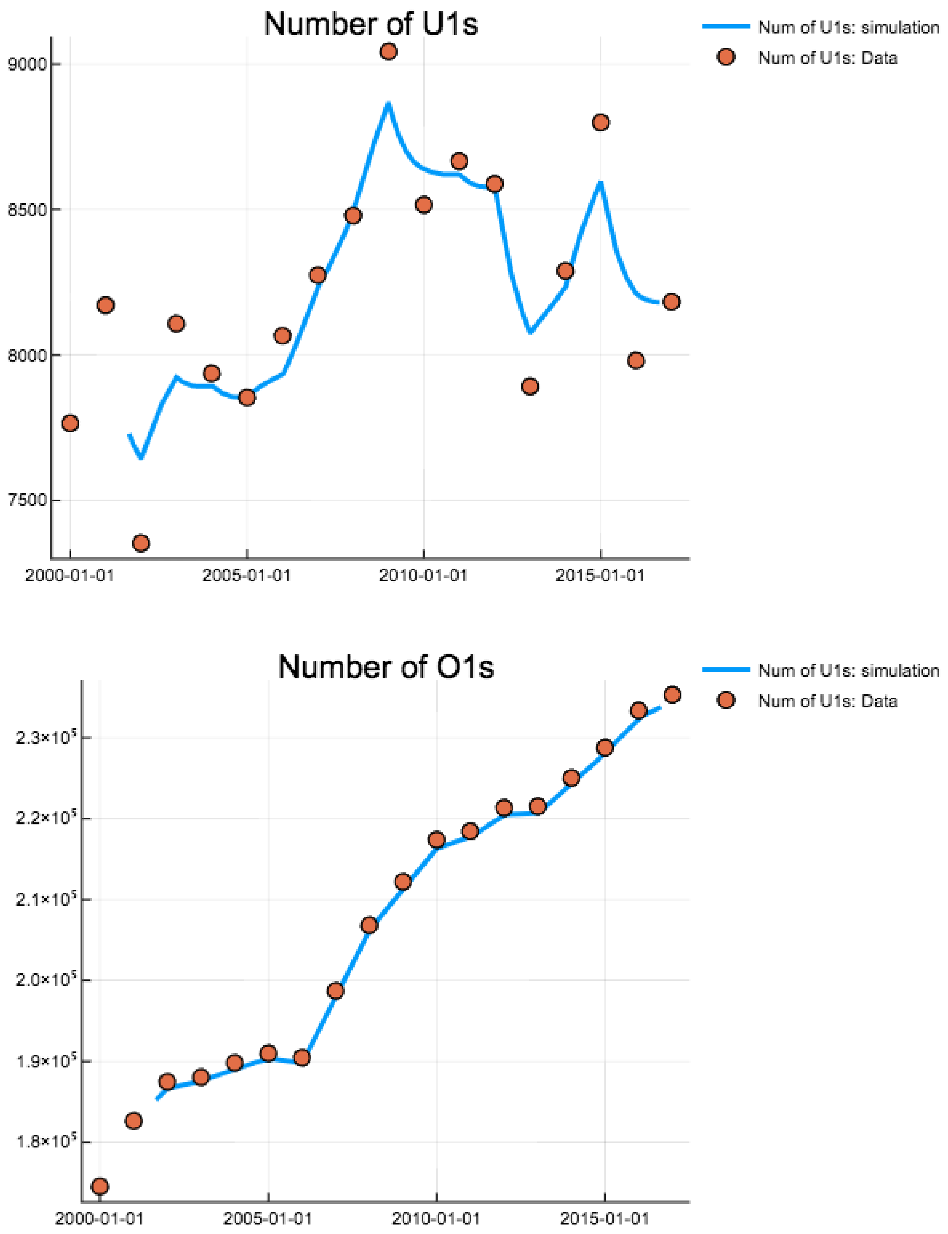


Figure 6: Comparison of total numbers of U1s and O1s on each 1st Jan 2000-2017 (dots) against simulated values (curve).

disease, multiplied by the probability that severely diseased individuals would require hospitalisation. The probability that an infected individual became diseased depended on whether it was the individual's primary infection episode or not. The underlying data for estimating these probabilities was drawn from cohort studies on RSV disease rates [13, 14]. We adapted these probabilities for our model using our assumption that all infected under-ones were experiencing their first RSV episode, and all over-ones were experiencing their second or subsequent infection.

- Inferred from the KCH hospitalisation data set (see below).

| Parameter | Description | Value | Data source |
|---|---|---|---|
| | **Parameters from literature** | | |
| $\sigma_{U1}$ | Susceptibility of under-ones | $1$ | [15] |
| $\sigma_{O1}$ | Susceptibility of over-ones | $0.75$ | [15] |
| $\nu$ | Rate of waning of immunity | 2 per year | [16] |
| $\gamma_1$ | Rate of recovery for under-ones | 1/9 per day | [17] |
| $\gamma_2$ | Rate of recovery for over-ones | 1/4 per day | [17] |
| $\iota_2$ | Factor reducing infectiousness for over-ones | 0.5 | [5] |
| | **Parameters estimated from KHDSS data** | | |
| $\mu(n,t)$ | Birth/turnover rate for households of size $n$ on day $t$ | Varies, see above | - |
| $r(n,t)$ | Rate of change of numbers of households of size $n$ on day $t$ | Varies, see above | - |
| $P_{H\to A,t}$ | Conditional age distribution given household config. on day $t$ | Varies, see above | - |
| $P_{A\to H,t}$ | Conditional household config. distribution given age category on day $t$ | Varies, see above | - |
| | **Parameters chosen by authors** | | |
| $\eta$ | Ageing rate for U1s | 1/365.25 per day | - |
| $\epsilon$ | Basic rate of external infections for whole population | 10 per day | - |

Table 1: Parameters from literature, estimated from KHDSS data or chosen by authors. Parameter estimates were originally gathered by Kinyanjui *et al*, and we have used our assumption that no susceptible U1s have been infected before and that all O1s have been infected at least once.

#### Parameter inference for the household- and age- model

As mentioned in the main text we used the EM algorithm [18] to estimate parameters for the model. Again, as described in the main text the parameters we chose for inference were:

- Infectious contact rate outside the household between U1s and all others in the community accessing KCH ($b_{U1}$).
- Infectious contact rate outside the household among all O1s in community ($b_{O1}$).
- Infectious contact rate within the household ($\tau$).
- Rate of loss of maternally derived immunity to RSV ($\alpha$).
- The joint normal distribution of the yearly log-seasonality amplitude and phase ($[\xi, \phi] \sim \mathcal{N}(\boldsymbol{\mu}, \boldsymbol{\Sigma})$).

Where the community age mixing matrix $T(a,b)$ was in block form:

$$T = \left(\begin{array}{c|c|c} b_{U1} & b_{U1} & b_{U1} \\ \hline b_{U1} & b_S + b_{O1} & b_{O1} \\ \hline b_{U1} & b_{O1} & b_{O1} \end{array}\right). \tag{38}$$

| Age category | Probability of hospitalisation per infection |
|---|---|
| 0-1 month | 0.10 |
| 1-2 month | 0.10 |
| 2-3 month | 0.063 |
| 3-4 month | 0.059 |
| 4-5 month | 0.054 |
| 5-6 month | 0.025 |
| 6-7 month | 0.019 |
| 7-8 month | 0.022 |
| 8-9 month | 0.012 |
| 9-10 month | 0.016 |
| 10-11 month | 0.013 |
| 11-12 month | $5.1\text{x}10^{-3}$ |
| 1-2 years old | $2.6\text{x}10^{-3}$ |
| 2-3 years old | $7.5\text{x}10^{-4}$ |
| 3-4 years old | $2.2\text{x}10^{-4}$ |
| 4-5 years old | $3.8\text{x}10^{-5}$ |

Table 2: Age-dependent hospitalisation probabilities per infection derived from Kinyanjui et al [5].

The log-likelihood for our model [equation (8) main text] was defined using the incidence rates $\mathcal{I}_a(t)$ predicted by solving the model. The incidence rate for all the households in the generic household configuration was,

$$\text{For U1s: } \mathcal{I}_h(U1,t) = (\sigma_{U1}\beta(t)s_1(\lambda_{hh} + \lambda_{com}(U1,h,t)))H_h(t) \tag{39}$$
$$\text{For O1s: } \mathcal{I}_h(O1,t) = (\sigma_{O1}\beta(t)s_2(\lambda_{hh} + \lambda_{com}(O1,h,t)))H_h(t). \tag{40}$$

Where the household force of infection for the generic household configuration was $\lambda_{hh} = \tau(i_1 + \iota_2 i_2)$. We converted the household incidence rate into an age structured incidence rate by using conditional age distributions, and this allowed us to calculate the cumulative hospitalisations in age category $a$, predicted by a given set of parameters and yearly seasonality realisations, in weekly intervals $w_i = (t_{i,1}, t_{i,2})$ using the age dependent hospitalisation rates per infection $h_a$ (see table 2)

$$\mathcal{I}_a(t) = \sum_{h\in\Sigma} \mathbb{P}(A \in a | M < A, A \leq 1 \text{ year})\mathcal{I}_h(U1,t) + \mathbb{P}(a|h, A > 1 \text{ year})\mathcal{I}_h(O1,t) \tag{41}$$

$$\mathcal{H}(a, w_i) = K(t)\int_{t_{i,1}}^{t_{i,2}} \mathcal{I}_a(t)h_a \,\mathrm{d}t, \tag{42}$$

$$\ln \mathbb{P}(\mathcal{D}_{i,a}|\theta, \boldsymbol{\xi}, \boldsymbol{\phi}) = l(\theta, \boldsymbol{\xi}, \boldsymbol{\phi}) = \sum_i \sum_a \ln f_{poi}(\mathcal{D}_{i,a}|\mathcal{H}(a, w_i)). \tag{43}$$

Here $K(t)$ is a time-varying scale factor that accounted for the fact that whilst we were modelling RSV infection for the KDHSS population, other individuals were accessing KCH for treatment of RSV-induced severe disease. To fit $K(t)$ we first performed a polynomial regression $R(t)$ against the ratio of KDHSS members using KCH against non-KDHSS members (figure 7)[2]. Having fitted the ratio, the scale factor was $K(t) = (1+R(t))/R(t)$, which we derived by assuming that non-residents were experiencing RSV hospitalisations at proportionally the same rate as residents.

[2] t = 0 (days) is 22nd April 2002 fitted curve is R(t) = 1.2409370942399072+ 0.002241544326043194 * $t$+ -2.4542080692642405e-6 * $t^2$+ 9.446287443394756e-10 * $t^3$+ -1.552135315860274e-13 * $t^4$ + 9.097449985032835e-18 * $t^5$. $R(t) = R(0)$ for $t < 0$, and $R(t)$ took its final value for times after 1st sept 2016.

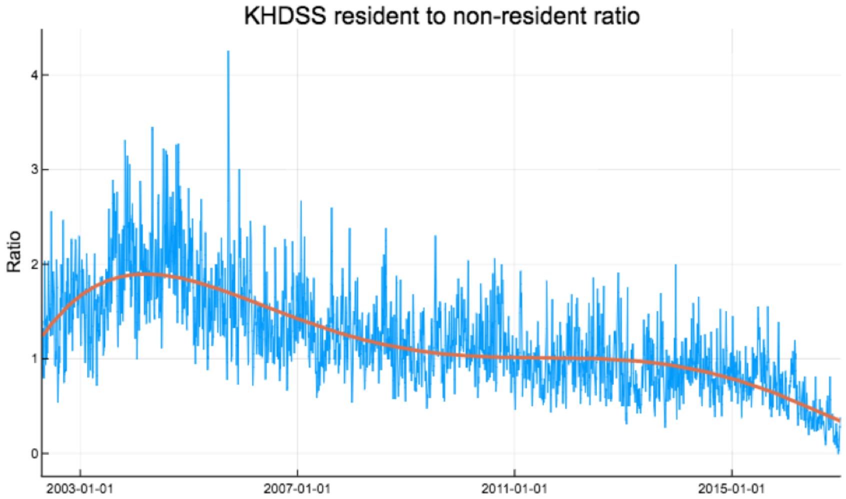


Figure 7: Ratio of KDHSS residents to non-residents weekly accessing KCH for confirmed RSV treatment. Red curve is polynomial fit $R(t)$.

The conditional age category of an U1 who has definitely been infected, where $a = (a_0, a_1)$,

$$\begin{aligned}\mathbb{P}(A \in a | M < A, A \leq 1 \text{ year}) &= \mathbf{1}(a \leq 1 \text{ year}) \frac{\mathbb{P}(M < A | A \in a)\mathbb{P}(A \in a | a \leq 1 \text{ year})}{\mathbb{P}(M < A | a \leq 1 \text{ year})} \\ &= \mathbf{1}(a \leq 1 \text{ year}) \frac{a_1 - a_0 + \overline{M}(e^{-a_1/\overline{M}} - e^{-a_0/\overline{M}})}{T(1 - e^{-T/\overline{M}})\sigma_{U1}} \end{aligned} \quad (44)$$

An implication of expression (44) is that if $a_0$ and $a_1$ are both significantly less than $\overline{M} = 1/\alpha$ then $\mathbb{P}(A \in a | M < A, A \leq 1 \text{ year}) \approx 0$; that is that, although we have assumed that the conditional age of an U1 is distributed evenly over the first year of life, the conditional age distribution of an U1 who has been infected is typically older than $\overline{M}$. This allowed us to extract information for inferring $\alpha$ from the age distribution of hospitalised children at KCH despite only using a crude U1/O1 age distinction in the mechanistic formulation of the household-and-age model. The log-likelihood $l(\theta, \boldsymbol{\xi}, \boldsymbol{\phi})$ [equation(43)] could be determined for a given set of parameters **and** realisations of the yearly seasonal amplitude and phase by solving the full ODE system numerically [equation (35)], and thereby also calculating the weekly hospitalisations. $\theta$ represented the model parameters to be inferred, $\boldsymbol{\xi}$ and $\boldsymbol{\phi}$ were the vectors of the seasonal transmission model equation (5), and $\mathcal{D}_i, a$ was the KCH hospitalisation data for the $i$th week in the $a$ age category.

The main difficulty in the inference for the unknown parameters $\theta$ was that the actual realisations of $\boldsymbol{\xi}$ and $\boldsymbol{\phi}$ are not observed, therefore $l(\theta, \boldsymbol{\xi}, \boldsymbol{\phi})$ could not be calculated directly. Instead, we use the EM algorithm to converge onto a maximiser of the marginal likelihood, $\mathcal{L}(\theta) = \int \mathbb{P}(\mathcal{D}, \boldsymbol{\xi}, \boldsymbol{\phi} | \theta) \, \mathrm{d}\boldsymbol{\xi} \, \mathrm{d}\boldsymbol{\phi}$. The EM algorithm converges a sequence of parameter estimates $(\theta^{(n)})_{n \geq 0}$ towards a local maximum of the marginal likelihood by alternatively, 1) calculating the expected value of the log-likelihood over the conditional distribution of $\boldsymbol{\xi}$ and $\boldsymbol{\phi}$ given the observed data $\mathcal{D}$ and the current estimate of the parameters, which we dub the $Q$ function [E step], and, 2) finding the parameters which maximised the $Q$ function [M step]. We now give details of how this was implemented for the specific model developed in this paper:

- E step: The conditional distribution of $\boldsymbol{\xi}$ and $\boldsymbol{\phi}$ given the $n$-th parameter estimate $\theta^{(n)}$, from the previous M-step, and $\mathcal{D}$ could not be calculated in closed form. In principle, this distribution could have estimated numerically (e.g. by using a particle filter method), however, because the household- and age-structured RSV transmission model was comparatively slow to integrate ($\sim$ 40 secs per simulation) we resorted to saddle-point integration. Our argument is that because nearly every year has a sharply peaked hospitalisation rate then, given a parameter estimate $\theta^{(n)}$, the conditional probability of $(\boldsymbol{\xi}, \boldsymbol{\phi})$ should be concentrated around a particular value, making saddle-point integration an appropriate approximation (see [19] for further details on saddle-point integration). Using the saddle-point approximation we could solve for the $Q$ function,

$$\begin{aligned} Q(\theta|\theta^{(n)}) &= \mathbb{E}_{\boldsymbol{\xi},\boldsymbol{\phi}|\mathcal{D},\theta^{(n)}}[\ln \mathbb{P}(\mathcal{D}, \boldsymbol{\xi}, \boldsymbol{\phi}|\theta)] \\ &= \mathbb{E}_{\boldsymbol{\xi},\boldsymbol{\phi}|\mathcal{D},\theta^{(n)}}[l(\theta, \boldsymbol{\xi}, \boldsymbol{\phi}) + \ln \mathbb{P}(\boldsymbol{\xi}, \boldsymbol{\phi}|\theta)] \\ &\approx l(\theta, \boldsymbol{\xi}^*, \boldsymbol{\phi}^*) + \ln \mathbb{P}(\boldsymbol{\xi}^*, \boldsymbol{\phi}^*|\theta) \\ &\propto l(\theta, \boldsymbol{\xi}^*, \boldsymbol{\phi}^*) - \sum_i [(\xi_i^* - m_\xi) \ (\phi_i^* - m_\phi)] \boldsymbol{\Sigma}_{\xi\phi}^{-1} [(\xi_i^* - m_\xi) \ (\phi_i^* - m_\phi)]^T. \end{aligned} \quad (45)$$

The approximation step in equation (45) is the saddle-point integration approximation of the average, and the quadratic form is due to our assumption that the seasonal amplitude and phases are distributed jointly normally. Saddle-point integration is equivalent to assuming that the full mass of the conditional distribution of $(\boldsymbol{\xi}, \boldsymbol{\phi})$ was concentrated at its most probable value,

$$\begin{aligned} (\boldsymbol{\xi}^*, \boldsymbol{\phi}^*) &= \arg\max_{\boldsymbol{\xi},\boldsymbol{\phi}} \ln \mathbb{P}(\boldsymbol{\xi}, \boldsymbol{\phi}|\mathcal{D}, \theta^{(n)}) \\ &= \arg\max_{\boldsymbol{\xi},\boldsymbol{\phi}} \{\ln \mathbb{P}(\mathcal{D}|\boldsymbol{\xi}, \boldsymbol{\phi}, \theta^{(n)}) + \ln \mathbb{P}(\boldsymbol{\xi}, \boldsymbol{\phi}|\theta^{(n)})\} \\ &= \arg\max_{\boldsymbol{\xi},\boldsymbol{\phi}} \{l(\theta^{(n)}, \boldsymbol{\xi}, \boldsymbol{\phi}) - \sum_i [(\xi_i^* - m_\xi^{(n)}) \ (\phi_i^* - m_\phi^{(n)})] \boldsymbol{\Sigma}_{\xi\phi}^{-1,(n)} [(\xi_i^* - m_\xi^{(n)}) \ (\phi_i^* - m_\phi^{(n)})]^T\}. \end{aligned} \quad (46)$$

We determined $(\boldsymbol{\xi}^*, \boldsymbol{\phi}^*)$ by sequentially optimising equation (46) over each season by simulating the model repeated and using the Nelder-Mead algorithm implemented within the **Optim** package for Julia 0.6. Note that saddle point integration has converted solving for the function $Q$ into a regularised maximum likelihood problem where the regularisation was provided by the mean and covariance matrix for log-seasonal amplitude and phase derived in the previous M step.

- M step: Having constructed the $Q$ function associated with the $n$-th parameter iteration [equation (45)], we maximised $Q$ over $\theta$. The maximum point of $Q$ being $\theta^{(n+1)}$ for the next E-step. Maximisation proceeded in three stages:

  1. The maximising values for the mean and covariance matrix of the random seasonal amplitude and phase were given by maximum likelihood using $(\boldsymbol{\xi}^*, \boldsymbol{\phi}^*)$ derived in the E-step. This was performed using the **fit_mle** function provided by the Julia **Distributions** package.
  2. We performed a global optimisation for $Q$ over a box in parameter space defined by limits $[0, 1]$ for transmission parameters and $1/\alpha = \overline{M} \in [10, 120]$ days for the inverse rate of loss of maternal immunity. Global optimisation was performed by running 600 iterations of a differential evolution optimiser [20] with 50 agents. The differential evolution optimiser was implemented by the **adaptive_de_rand_1_bin_radiuslimited** optimiser from the Julia **BlackBoxOptim** package. The purpose of the global optimisation step was to reduce the dependence on choosing an initial guess about $\theta$ since the whole plausibility space of the

parameters was explored at each iteration of the EM algorithm. We called the best performing agent's parameter set on the $(n+1)$th step, $\tilde{\theta}^{(n+1)}$.

3. We used $\tilde{\theta}^{(n+1)}$ as the starting point for a further local optimisation of $Q$ using the Nelder-Mead algorithm implemented by the Julia **Optim** package. This step provided $\theta^{(n+1)}$ for the next E-step.

We iterated EM algorithm until no further improvement in the value of $Q^* = \max_\theta Q$ was achieved, and then retained $\theta^* = \arg\max_\theta Q$ as the maximum likelihood estimator for the parameters. 95% confidence intervals were estimated by using univariate profile likelihood for $Q$; that is varying one parameter at a time whilst keeping others fixed until a $\chi^2$ region was determined around the maximum of $Q$ (see King *et al* for a description of 95% CIs for dynamical systems [21]).

#### School mixing scenarios and inference results

We were unable to identify a mixing rate within schools $b_S$, see equation (38), therefore we considered four values of $b_S$ each determined by what a baseline reproductive value for RSV would be *if* only school children mixed together and the seasonality was just $\beta(t) = 1$, $R_S$, using the simple formula,

$$R_S = \frac{b_S \sigma_{O1} \iota_2}{\gamma_2} \tag{47}$$

These four scenarios were: zero schools transmission ($R_S = 0$), low schools transmission ($R_S = 0.5$), medium schools transmission ($R_S = 1$), and, high schools transmission ($R_S = 1.5$). We saw that once maximum likelihood estimation was performed on the free parameters: $\theta = (b_{U1}, b_{O1}, \tau, \alpha, \boldsymbol{m}, \Sigma_{\xi\phi})$ the resultant fits to the data were very similar visually (see figure 8). We noticed that the outcomes of vaccination were also similar for each four scenarios (see below and figure 10). Therefore, for robustness of conclusion we used the most pessimistic scenario within the main body of the paper, which was high schools transmission $R_S = 1.5$. The maximum likelihood estimates for parameters using the high schools transmission scenario are given in table 3, and the maximum likelihood estimates for all scenarios summarised in figure 9.

| Parameter | Description | Value |
|---|---|---|
| $b_{U1}$ | Community transmission rate for U1s | 0.22 [0.18,0.27] per day |
| $b_{O1}$ | Community transmission rate for O1s | 0.20 [0.18,0.21] per day |
| $\tau$ | Transmission rate to *each* other member of household | 0.040 [0.032, 0.048] per day |
| $\overline{M}$ | Mean duration of maternal protection at birth | 21.6 [17.2, 26.1] days |
| $m_\xi$ | Mean amplitude of log-seasonality | 0.61 [0.51, 0.72] |
| $m_\phi$ | Mean timing of log-seasonality peak (phase) | 67.7 [40.2, 77.7] days |
| $\sigma_\xi$ | Std. amplitude of log-seasonality | 0.20 [0.098,0.31] |
| $\sigma_\phi$ | Std. timing of log-seasonality peak (phase) | 38.7 [30.0, 48.5] days |
| $\rho_{\xi\phi}$ | Corr. coefficient between log-seasonal amplitude and phase | -0.035 [-0.12, 0.072] |

Table 3: Parameters from literature, estimated from KHDSS data or chosen by authors. Parameter estimates were originally gathered by Kinyanjui *et al*, and we have used our assumption that no susceptible U1s have been infected before and that all O1s have been infected at least once.

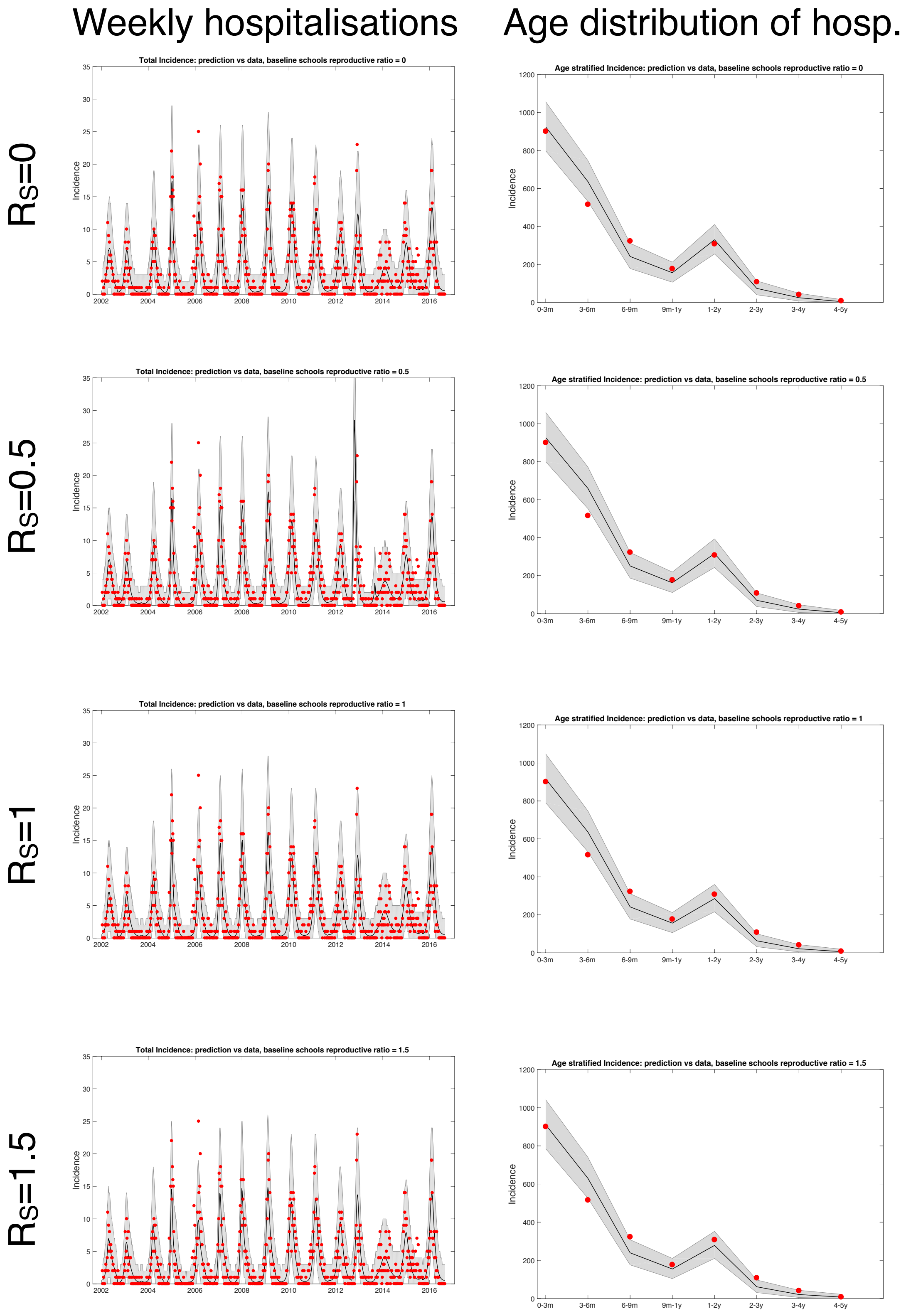


Figure 8: Plots of fitted weekly hospitalisations and the age distribution of hospitalisations for four scenarios (differing values of the schools based baseline $R_S$). In each case, parameter inference was performed and the maximum likelihood estimators used.

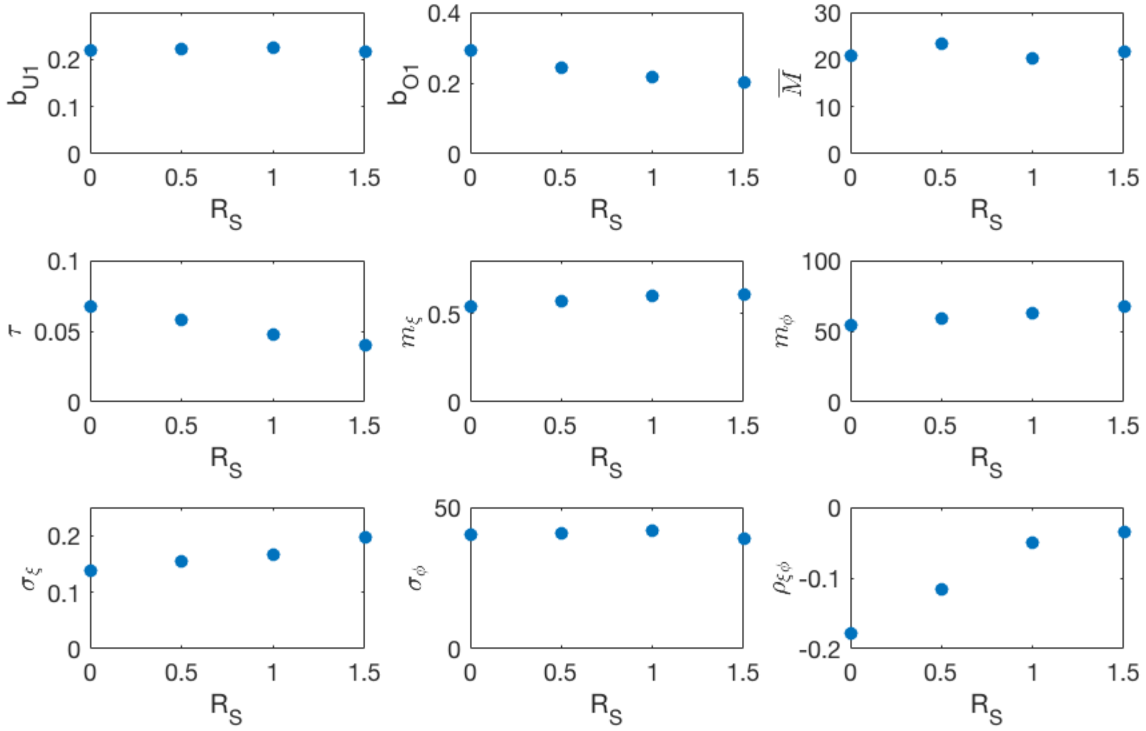


Figure 9: Maximum likelihood parameters for the different school transmission rate scenarios. $b_{U1}$, $b_{O1}$ are respectively the under-one and over-one mixing components of the community mixing rate matrix. $\tau$ is the rate at which a household member infectiously contacts *each* other household member. $\overline{M} = 1/\alpha$ is the mean period of maternal protection after birth. $\boldsymbol{m} = (m_\xi \; m_\phi)$ is the mean vector of the random seasonality, and $\sigma_\xi$, $\sigma_\phi$ and $\rho_{\xi\phi}$ are respectively the standard deviations of the seasonal amplitude, seasonal phase and the correlation between the two, derived from the estimated covariance matrix $\Sigma_{\xi\phi}$.

#### Modelling vaccination in the household- and age-structured RSV transmission model

As described in the main paper we modelled the use of two different vaccines: a vaccine deployed to boost the period during which a newborn is protected from RSV by an unknown period $P$ with coverage $V_{cov}$ [MAB vaccine], and a vaccine deployed to O1 household members of the newborn which provokes a period of protection to RSV infection similar to the immunity period of a natural infection at household coverage $H_{cov}$ [IRP vaccine]. Already infected or recovered O1s were not affected by the IRP vaccine. We assumed that the MAB and IRP vaccines were deployed independently, which is useful for gauging potential effectiveness, but unrealistic. In reality, any reason a mother-to-be might miss being MAB vaccinated would also be a reason that the household O1s wouldn't get vaccinated.

The IRP vaccine altered the effective birth events by also provoking transitions to $R_2$ state at the point of birth,

- <u>Demographic turnover due to births with vaccination:</u>

$$[s_1, i_1, r_1, s_2, i_2, r_2] \to [s_1 + 1, i_1, r_1, s_2 - 1, i_2, r_2] \text{ at rate: } (1 - H_{cov})\mu(n,t)s_2, \quad (48)$$
$$[s_1, i_1, r_1, s_2, i_2, r_2] \to [s_1 + 1, i_1, r_1, s_2, i_2 - 1, r_2] \text{ at rate: } (1 - H_{cov})\mu(n,t)i_2, \quad (49)$$
$$[s_1, i_1, r_1, s_2, i_2, r_2] \to [s_1 + 1, i_1, r_1, s_2, i_2, r_2 - 1] \text{ at rate: } (1 - H_{cov})\mu(n,t)r_2, \quad (50)$$
$$[s_1, i_1, r_1, s_2, i_2, r_2] \to [s_1 + 1, i_1, r_1, 0, i_2, s_2 + r_2 - 1] \text{ at rate: } H_{cov}\mu(n,t)(s_2 + r_2), \quad (51)$$
$$[s_1, i_1, r_1, s_2, i_2, r_2] \to [s_1 + 1, i_1, r_1, 0, i_2 - 1, s_2 + r_2] \text{ at rate: } H_{cov}\mu(n,t)i_2. \quad (52)$$

The MAB vaccine altered both the probability that an U1 is protected, and the age distribution of those who are infected. We denote the random period of time a newborn born to a MAB vaccinated mother is protected from RSV as $M_{vac} = M + P$, which has distribution function,

$$\mathbb{P}(M_{vac} \leq a) = \begin{cases} 0 & 0 \leq a \leq P \\ (1 - \exp(-(a-P)/\bar{M}))/(1 - \exp(-(T-P)/\bar{M})) & P \leq a \leq T \\ 1 & \text{otherwise} \end{cases} \quad (53)$$

The mean susceptibility of U1s after MAB vaccination has been applied to the population was,

$$\begin{aligned} \sigma_{U1,vac} &= \frac{1}{T}\int_0^T \big((1 - V_{cov})\mathbb{P}(M \leq a) + V_{cov}\mathbb{P}(M_{vac} \leq a)\big)\,\mathrm{d}a \\ &= 1 - \frac{\overline{M}}{T} + (1 - V_{cov})P\frac{e^{-T/\overline{M}}}{1 - e^{-T/\overline{M}}} + V_{cov}\frac{(T-P)e^{-(T-P)/\overline{M}}}{T(1 - e^{-(T-P)/\overline{M}})} - V_{cov}\frac{P}{T}. \end{aligned} \quad (54)$$

The conditional age category of an U1 who has definitely been infected, where $a = (a_0, a_1)$, after MAB vaccine has been deployed at coverage $V_{cov}$ was,

$$\begin{aligned} \mathbb{P}(A \in a | \tilde{M} < A, A \leq 1 \text{ year}) &= \mathbf{1}(a \leq 1 \text{ year})\frac{\big((1 - V_{cov})\mathbb{P}(M < A | A \in a) + V_{cov}\mathbb{P}(M_{vac} < A | A \in a)\big)\mathbb{P}(A \in a | a \leq 1 \text{ year})}{\mathbb{P}(M < A | a \leq 1 \text{ year})} \\ &= \frac{\mathbf{1}(a \leq 1 \text{ year})}{T\sigma_{U1,vac}}\Big((1 - V_{cov})\frac{a_1 - a_0 + \overline{M}(e^{-a_1/\overline{M}} - e^{-a_0/\overline{M}})}{1 - e^{-T/\overline{M}}} + V_{cov}f(a,P)\Big). \end{aligned} \quad (55)$$

Where $\tilde{M}$ is the random maternal protection duration of a newborn before we observe whether the newborn's mother had been MAB vaccinated. The function $f(a, P)$ completes equation (55) by giving the age distribution of U1s who had boosted maternal protection to RSV but was nonetheless infected,

$$f(a,P) = \begin{cases} 0 & a_0 \leq P \text{ and } a_1 \leq P \\ \frac{a_1 - P + \overline{M}(e^{-(a_1 - P)/\overline{M}} - 1)}{1 - e^{-(T-S)/\bar{M}}} & a_0 \leq P \text{ and } a_1 > P \\ \frac{a_1 - a_0 + \overline{M}(e^{-(a_1 - P)/\overline{M}} - e^{-(a_0 - P)/\overline{M}})}{1 - e^{-(T-S)/\bar{M}}} & a_0 > P \text{ and } a_1 > P \end{cases} \quad (56)$$

Note that because $\sigma_{U1,vac}$ depended on $V_{cov}$ the age distribution of infected U1s depended on $V_{cov}$ in a nonlinear fashion.

We considered a range of values for $P$ and $H_{cov}$ for each of the schools transmission scenarios; using the maximum likelihood estimators for the inferred parameters for each scenario. In each scenario, at $V_{cov} = 1$ the median reduction in hospitalisations was similar, although for the high school transmission scenario vaccination was slightly less effective (figure 10). Therefore, we used this scenario in the main paper as a pessimistic/robust example. As mentioned in main text we simulated 10 years into the future over 500 independent realisations of the random seasonality. Presented are medians of % reduction in hospitalisations at KCH compared to no intervention.

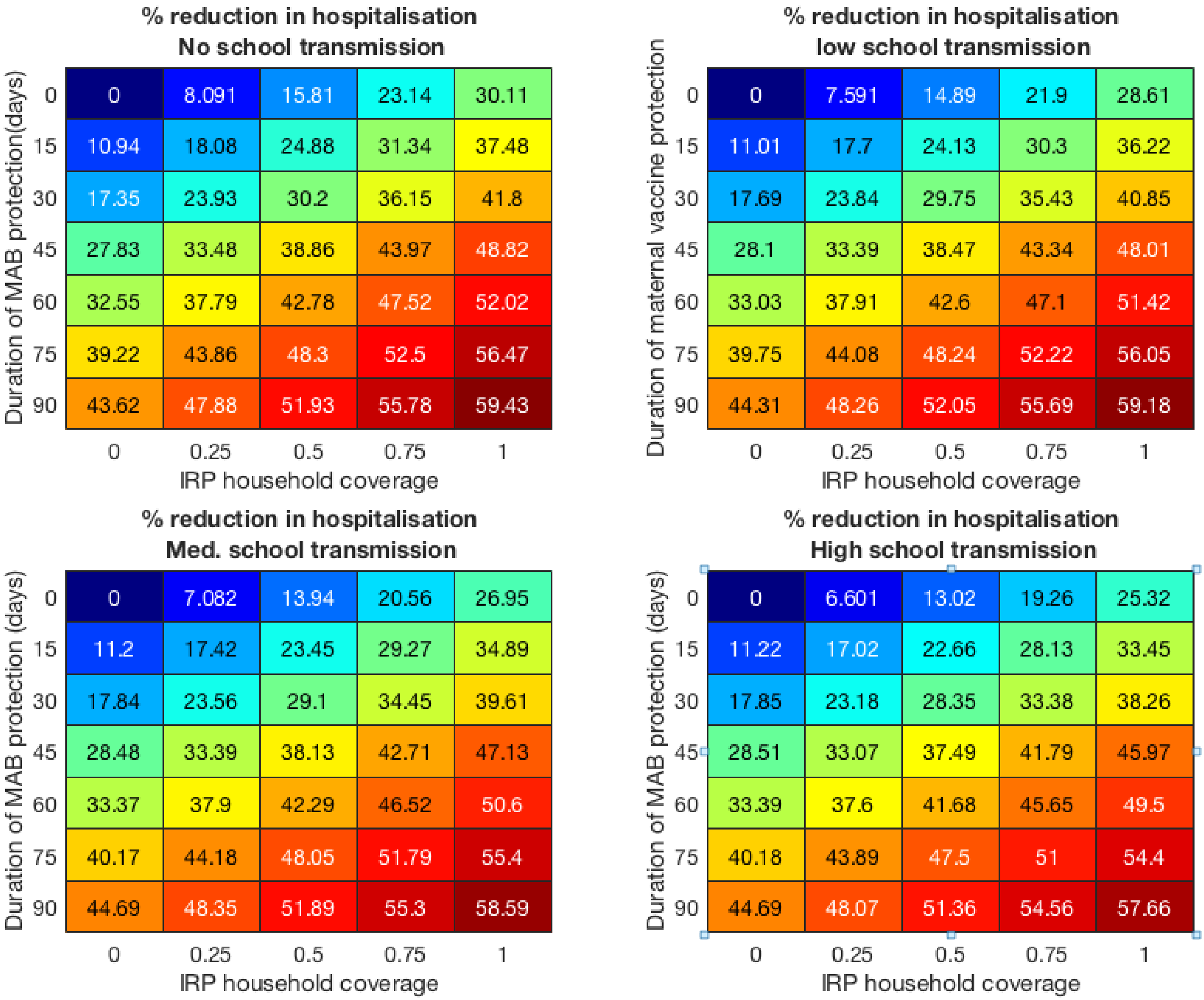


Figure 10: Vaccine effectiveness for the four school mixing scenarios at 100% MAB coverage.
